# MRTF-dependent cytoskeletal dynamics drive efficient cell cycle progression

**DOI:** 10.1101/2025.06.06.657372

**Authors:** Julie C. Nielsen, Maria Benito-Jardon, Noel Christo Petrela, Jessica Diring, Sofie Bellamy, Richard Treisman

## Abstract

Serum response factor (SRF) and its cofactors, Myocardin-related transcription factors A/B (MRTF-A/B), regulate transcription of numerous cytoskeletal structural and regulatory genes, and most MRTF/SRF inactivation phenotypes reflect deficits in cytoskeletal dynamics. We show that MRTF-SRF activity is required for effective proliferation of both primary and immortalised fibroblast and epithelial cells. Cells lacking the MRTFs or SRF proliferate very slowly, express elevated levels of SASP factors and SA-β-galactosidase activity, and inhibit proliferation of co-cultured primary wildtype cells. They exhibit decreased levels of CDK1 and CKS2 proteins, and elevated levels of CDK inhibitors, usually CDKN1B/p27. These phenotypes, which can be fully reversed by re-expression of MRTF-A, are also seen in wildtype cells arrested by serum deprivation. Moreover, in wildtype cells direct interference with cytoskeletal dynamics through inhibition of ROCKs or Myosin ATPase induces a similar proliferative defect to that seen in MRTF-null cells. MRTF-null cells exhibit multiple cytoskeletal defects, and markedly reduced contractility. We propose that MRTF-SRF driven cytoskeletal dynamics and contractility are essential for operation of the pro-proliferative signal provided by cell-substrate adhesion.

## INTRODUCTION

The cell cycle is controlled by a conserved network of Cyclin/CDKs and their regulators that ensures that cell growth, genome replication and segregation, and cell division occur in the right order and at the right time (Malumbres & Barbacid, 2001; Morgan, 1997). In multicellular organisms, however, many cell types remain quiescent unless prompted to proliferate by external cues (Marescal & Cheeseman, 2020). Proliferation of cultured mammalian cells is thus dependent on external stimuli, including serum mitogens, nutrients, and substrate adhesion, whose withdrawal or downregulation by other cues, such as cell confluence and matrix compliance, leads to cell cycle exit and return to quiescence (Otsuka & Moskowitz, 1975; Pardee, 1974; Pardee *et al*, 1978). These requirements and controls are relaxed in cultured tumour cells, suggesting that their disruption is important for oncogenic transformation (Malumbres & Barbacid, 2001). Stress stimuli such as telomere erosion or oncogene activation lead to senescence, a distinct non-proliferative state considered irreversible (Campisi & d’Adda di Fagagna, 2007; Salama *et al*, 2014). Senescent cells secrete factors that promote cell cycle arrest in both an autocrine and paracrine manner (Acosta *et al*, 2013; Coppe *et al*, 2008).

In non-transformed cells, the Ras-ERK and Rho GTPase signalling pathways are central to control activity of the core cell cycle machinery by mitogenic stimuli (Pruitt & Der, 2001) (Olson *et al*, 1995). Ras-ERK signalling is linked to expression of Cyclin D mRNA and protein synthesis (Matsushime *et al*, 1991; Won *et al*, 1992), while Rho signalling controls focal adhesion assembly and signalling, and promotes actin-dependent cell contractility (Hotchin, 1995 #52, reviewed by Burridge & Chrzanowska-Wodnicka, 1996). Proliferation requires Rho-dependent adhesion and cell spreading (Chen, 1997 #55, reviewed by Kamranvar *et al*, 2022; Mammoto & Ingber, 2009), and Rho-regulated cytoskeletal dynamics play critical roles in mitosis and cytokinesis (Basant & Glotzer, 2018; Ramkumar & Baum, 2016). Classical studies showed that cell cycle re-entry from quiescence requires new RNA synthesis by pre-existing cellular factors (Pledger *et al*, 1981; Smith & Stiles, 1981). Mitogen-induced cellular immediate-early (IE) genes include c-myc and c-fos, other transcription factors, and actins (Cochran *et al*, 1983; Cochran *et al*, 1984; Greenberg & Ziff, 1984; Kelly *et al*, 1983), and IE genes are also induced upon cell-ECM adhesion (Dike & Farmer, 1988). Nevertheless, whether and how IE gene induction controls activation of cyclin D remains unclear.

Many IE genes are controlled by the SRF transcription factor (Norman *et al*, 1988; Schratt *et al*, 2001; Treisman, 1986). SRF works in partnership with two families of signal-regulated co-factors, the TCFs and the MRTFs, which respectively couple its activity to the Ras-ERK and Rho signalling pathways (Olson & Nordheim, 2010; Posern & Treisman, 2006), MRTF-SRF target genes include actins and many other cytoskeletal structural and regulatory proteins (Esnault *et al*, 2014; Olson & Nordheim, 2010), and cells lacking SRF or the MRTFs exhibit deficits in adhesion, motility and invasion (Alberti *et al*, 2005; Medjkane *et al*, 2009; Mokalled *et al*, 2010; Schratt *et al*, 2002). SRF-null mouse embryonic stem cells proliferate normally, as do embryos up to E6 (Arsenian *et al*, 1998; Schratt *et al*., 2001) and cell types such as neuroblasts (Alberti *et al*., 2005; Medjkane *et al*., 2009; Mokalled *et al*., 2010; Schratt *et al*., 2002), pre-DP thymocytes (Mylona *et al*, 2011) and TCR-activated T cells (Maurice *et al*, 2024). In contrast, SRF inactivation induces apparent senescence in smooth muscle cells (Angstenberger *et al*, 2007; Werth *et al*, 2010), and activates oncogene-induced senescence in cancer cells lacking the DLC1 tumour suppressor, a RhoGAP (Hampl *et al*, 2013; Hermanns *et al*, 2017; Muehlich *et al*, 2012).

Here we revisit the issue of Rho-dependent cytoskeletal dynamics, MRTF-SRF activity, and cell proliferation. We show that primary and immortalised fibroblasts and epithelial cells exhibit a proliferation defect with features of senescence and quiescence, but which can be reversed by MRTF-A re-expression. A similar proliferation defect is induced in wildtype MEFs upon serum starvation or stringent inhibition of ROCKs or myosin ATPase, and MRTF-null cells exhibit cytoskeletal defects and reduced contractility. Our results support a model in which MRTF-SRF linked cytoskeletal dynamics, particularly in cell adhesion and contractility, are essential for generation of the proliferative signal provided by cell adhesion.

## MATERIALS AND METHODS

### Mouse embryo fibroblast culture

Animals used were held under UK Home Office project licence P0389970. Primary fibroblasts were obtained from lung or kidney tissue from WT (*Gt(ROSA)26Sor^tm9(cre/ESR1)Arte^*) or *Mrtfa^-/-^*;*Mrtfb^fl/fl^* (*Mrtfa^tm1Eno^*;*Mrtfb^tm2.1Eno^*; *Gt(ROSA)26Sor^tm9(cre/ESR1)Arte^*) mice after tissue dissociation with Liberase TM and TH (Roche, 43739557/43740136). Primary fibroblasts and SV40 immortalised MEFs were maintained in DMEM with 10% foetal calf serum (FCS) and 1% Pen/Strep. Where indicated, cells were starved for 5 days in 0.3% FCS. For all experiments with primary fibroblasts we included biological replicates, meaning cells isolated from three different mice for each genotype. For MEFs, cells were isolated from three individual embryos per genotype, yielding three WT MEF pools (79, 80, and 82) and three *Mrtfa^-/-^*;*Mrtfb^fl/fl^* MEF pools (84, 86, and 89), as well as three SRF WT (MEF1, MEF3, and MEF4) and three *Srf^fl/fl^* MEF pools (Bis1, 2+, and Bis3 derived from *Srf^tm1Zli^;Gt(Rosa)26Sor^tm9(cre/ESR1)Arte^*). For all cell types (MEFs, primary fibroblasts and epithelial progenitor cells), *Mrtfb* inactivation was induced by addition of 1 µM 4-Hydroxytamoxifen (4OHT). For growth curves, 10,000 cells were seeded. Cells were treated with 5 µM H1152 or 10 µM blebbistatin as indicated.

### Retroviral transduction

Cell lines expressing Dox-inducible genes were constructed by lentiviral transduction of the *Mrtfa^-/-^Mrtfb^fl/fl^* MEF pool 86. Lentiviruses used were derivatives of pCW-MRTF-A-HA, which encodes Doxycycline-inducible C-terminally HA-tagged full-length mouse MRTF-A, together with IRES-expressed GFP, derived from pCW (Addgene #41393). For inducible expression of Rho-A^G14V^, CDK1, CKS2, or myoferlin, the MRTF-A sequences were substituted with appropriate cDNA. For lentivirus production Phoenix cells were transfected with pCW derivatives, pxPAX2 (packaging), and VSVG (envelope) in ratio 10:5:1 using Fugene HD (E2311, Promega) in OptiMEM (Gibco) with a Fugene:DNA ratio of 7:1, and selected using 1.25µg/mL puromycin. Cells were used as pools, except for *Mrtfa^-/-^Mrtfb^fl/fl^*DoxMRTF-A-HA cells, for which three cloned lines (1F2, 1F5, 1B2) were analysed. Protein expression was induced with 1µg/mL doxycycline.

### CRISPR/Cas9 editing

Dox-inducible Cas9 was inserted into the mH11 locus of *Mrtfa^-/-^*;*Mrtfb^fl/fl^*MEF pool 86 using targeted CRISPR (SH100+SH305 targeting mH11, GeneCopoeia). Following selection in 2µg/mL blasticidin, cloned cells were assayed for Dox-specific Cas9 editing, and line 2H17 was selected on the basis of growth behaviour comparable to the parental 86 pool (proliferation, SA-βGal, cell cycle protein expression, SASP mRNA expression, with and without 4OHT). For gene inactivation, 2H17 cells were infected with lentivirus expressing p27 sgRNA (Table S1), and selected with 1.25 µg/mL puromycin. Cas9 expression was induced with 1 µg/mL Doxycycline and three clonal lines – 3A2, 1F3 and 3B10 – of p27^-/-^ *Mrtfa^-/-^*;*Mrtfb^fl/fl^* MEFs isolated. Inactivation of p27 was confirmed by immunoblotting and sequencing (KO scores >90%; Synthego, ICE analysis).

p27^-/-^ *Mrtfa^-/-^*;*Mrtfb^fl/fl^* line 3A2 was then used to inactivate p21 and p57 using synthetic sgRNAs (Table S1), using RNAiMAX for sgRNA delivery. Following induction of Cas9 expression, three lines of p27^-/-^ p21^-/-^ p57^-/-^ *Mrtfa^-/-^*;*Mrtfb^fl/fl^* MEFs (3.1D10, 3.1C11, 3.1G9) were cloned and p21 and p57 inactivation confirmed by sequencing and immunoblotting. (KO scores >90% and >80%; Synthego, ICE analysis). For p57, sanger sequencing clone 3.1D10 revealed a 2bp deletion in exon 2 (5’ CGAGACGG 3’ -> 5’ CGA/CGG 3’).

### Epithelial cell cultures

Tracheal epithelial cultures were generated as described (Crotta *et al*, 2023; You *et al*, 2002). 4 to 6 tracheas from 10-20 week-old female mice were dissected, pooled, cut in pieces and digested overnight at 4°C with 3 mg/mL Pronase (Roche, 11459643001) in MTEC basic medium (15 mM HEPES (Gibco, 83264), 0.03% NaHCO3 (Gibco, 25080), 100 U/mL Penicillin, 0.1 mg/mL Streptomycin (Sigma, P4333), 2 mM L-Glutamine (Gibco, 25030081) in DMEM/F12 medium (Gibco, 21331020)). Cells (without cartilage) were washed with MTEC basic medium, and treated with 2 mg/mL DNase (Sigma, D4527) in MTEC basic medium for 10 minutes at room temperature. Cells were washed with MTEC basic medium and resuspended in MTEC plus (MTEC containing 25 ng/mL EGF (BD, 354001), 0.1 mg/mL D-valine (Sigma, V1255), 30 μg/mL Bovine Pituitary Extract (Gibco, 13028- 014), 0.1 μg/mL Cholera Toxin (Sigma, C8052), 1x ITS (Insulin, Transferrin, Selenium; Gibco, 41400045), 0.01 μM Retinoic Acid (Sigma, R2625), 10 μM Y27632; (Sigma, Y0503), 250 ng/mL Amphotericin B (Gibco, A2942), 10% FCS). After plating on an uncoated flask for 4 hours to allow fibroblasts to adhere, the epithelial cell supernatant was plated on 0.4 μg/cm^2^ fibronectin (BD, 356008), 4 μg/cm^2^ collagen (BD, 354236), treated with 4OHT the next day and cultured for 7 days until confluent. Cells were trypsinised (0.05% Trypsin, 30 min) (Gibco, 25300), resuspended in MTEC plus medium without Y27632 and seeded in non-differentiating conditions at 10000 cells per insert on clear polyester membrane inserts (0.4 μm pores; Sarstedt, 833932041) coated with 0.4 μg/cm^2^ fibronectin and 4 μg/cm^2^ collagen in a 24-well plate.

### Adipocyte differentiation

MEFs were cultured for 20 days in DMEM (10% FCS and 1% P/S) containing 0.5 mM IBMX, 1 µM dexamethasone, and 10 µM insulin without passaging. Cells were fixed in 10% neutral buffered formalin (NBF), incubated in 60% isopropanol, stained with 1.8 mg/mL Oil Red O, 66% isopropanol, and rinsed in water. Imaging used a Zeiss Axio Observer Z1 inverted microscope equipped with a QImaging colour camera.

### Immunoblotting

Cells were lysed in RIPA buffer (20 mM Tris-HCl pH 7.4, 150 mM NaCl, 0.1% sodium dodecyl sulfate (SDS), 0.5% Na-deoxycholate, 1% Triton X-100, 1x complete EDTA-free protease inhibitor cocktail (Roche, 54925800), 5 mM sodium fluoride, and 1 mM sodium orthovanadate) and after spinning down cell debris, protein concentration was normalised using a Bradford Assay (BioRad, 5000006). After addition of Laemmli Sample Buffer, samples were run at 120V on a NuPAGE 4-12% Bis-Tris gel (Invitrogen, NP0321BOX) in MES running buffer (Invitrogen, NP0002) and transferred to a nitrocellulose membrane (Amersham, 10600003) in transfer buffer (10% methanol, 192 mM glycine, 25 mM Trizma Base) at 200 mA for 1.5 hours. Membranes were incubated in primary antibodies (Table S2) overnight followed by secondary antibodies (IRDyes, LICOR; Table S2) and developed using Odyssey CLx (LICOR).

### Immunofluorescence and microscopy

Antibodies used are listed in Table S2. Cells were seeded on coverslips, fixed in 4% paraformaldehyde (Thermo Fisher Scientific, J19943.K2), permeabilised in 0.01% Triton X-100, blocked in 3% BSA and stained with primary antibodies at 4°C overnight. Coverslips were incubated with secondary antibodies in combination with DAPI and phalloidin-TRITC/647nm (Sigma, P1951/Invitrogen, A22287), where indicated, and finally mounted in mowiol mounting medium (9.32% (w/v) mowiol 4- 88, Calbiochem 475904). Images were acquired with the ZEN 2.3 SP1software using a Zeiss LSM 710 confocal microscope equipped with an AxioCam camera or a Yokogawa CSU-W1 SoRa Spinning Disk system with a Nikon Ti2 inverted microscope equipped with a BSI express camera (Teledyne Photometrics) using the NIS-Elements software (Nikon Instruments, Inc.). Images were processed using ImageJ.

For adhesion and spreading analysis, cells were incubated with CellMask-Orange (Thermo Fisher Scientific, C10045) 1:1000 in growth medium for 10 minutes followed by fixation and mounting as above. Imaging used a Zeiss Microscope AXIO Observer.D1 equipped with an AxioCam MRm camera, using the ZEN 2012 (blue edition) software. Area and circularity were measured using ImageJ.

For Phasefocus livecyte microscopy, cells were sparsely seeded in 12- or 24- well black glass-bottom plates. Images were acquired using the 10x objective and three large scan regions per well. Analysis used the Phasefocus analysis software.

### F-G actin fractionation

Cells were lysed in F-actin stabilisation buffer (100 mM PIPES pH 6.9, 5 mM MgCl_2_, 1 mM EGTA, 30% glycerol, 0.1% Triton x-100, 0.1% NP-40, 0.1% Tween, 0.15% β-mercaptoethanol, 1 mM ATP and 1x EDTA-free protease inhibitors) and homogenised using a 25G needle at 37°C. F- and G-actin fractions were obtained by ultracentrifugation at 100,000 xg for 1 hour at 37°C.

### Cell cycle phase, DNA synthesis and SA-βGal assays

Cells were pulsed with BrdU 2 hours pulse with BrdU, fixed in 100% ethanol on ice and treated with 2M HCl for 30 minutes. Cells were stained using anti-BrdU antibody (BD Biosciences, 347580) and resuspended in 50 µL 100 µg/mL ribonuclease A (Sigma, R5125) and 150 µL 50 µg/mL propidium iodide (PI, Sigma, P4170) before analysis by flow cytometry.

For EdU incorporation assays, cells were pulsed for 6h with Click-iT EdU with Alexa fluor 488 (C10337, Invitrogen), fixed in 3.7% PFA and permeabilised in 0.5% Triton X-100, and incubated in the Click-iT reaction cocktail and imaged after Hoechst staining and mounting in mowiol. Coverslips with micropatterns and micropillars were imaged using an Evident/Olympus VS200 slide scanner. Images were analysed in QuPath; EdU-positive cells were quantified using a threshold defined from a no EdU control, and Hoechst staining for total cell number.

SA-βGal activity analysis used the Senescence Cells Histochemical Staining Kit (Sigma, CS0030) and imaged on a Zeiss Axio Observer Z1 inverted microscope equipped with a QImaging colour camera. For flow-cytometric analysis of SA-βGal, cells were fixed in 2% PFA, washed in 1% BSA in PBS, and stained overnight with CellEvent Senescence Green (Thermo Fisher, C10841) according to the manufacturer’s protocol before analysis by flow cytometry.

For induction of proliferation phenotypes in wildtype cells by co-culture, primary kidney fibroblasts were infected with lentiviral particles containing pLVX-mCherry (Addgene #180646) two days after isolation, with 1.25 µg/mL puromycin selection for five days. Cells were then seeded either alone or in 100:1 co-culture with WT or *Mrtfab*-null MEFs for five days before CellEvent Senescence Green staining, and analysis by flow cytometry.

Apoptosis was measured using a luminogenic caspase-3/7 substrate from ApoTox-Glo Triplex Assay (G6320, Promega).

### RT-qPCR

RNA was extracted using GenElute Mammalian Total RNA miniprep kit (Sigma, RTN350-1KT) and cDNA synthesised using the Transcriptor First Strand cDNA Synthesis System (Roche, 0489703001). The cDNA was loaded into a MicroAmp Optical 384 well plate (Applied Biosystems, 4309849) and qPCR was carried out using 2x PowerUp SYBR Green Master Mix (Applied Biosystems, A25742). Absolute quantification of cDNA abundance was calculated using a mouse genomic DNA standard curve and then normalised to *Gapdh* or *Rps16* abundance. Datapoints are means of triplicate determinations. Primers used for RT-qPCR are listed in Table S1.

### RNA-Seq

RNA was isolated as above and libraries were prepared using PolyA KAPA mRNA HyperPrep kit (Roche, KK8581) and sequenced with single-end read mode on a HiSeq Illumina platform. The analysis was carried out with nf-core/rnaseq v3.5 using the GRCm38 as reference genome. Differential expression analysis was carried out with DEseq2. Following shrinking by the “ashr” method, log2 fold-change was plotted against the −log10 p-value to generate volcano plots. P-values were adjusted using the false discovery rate correction for multiple testing and Gene Ontology and Gene Set Enrichment Analysis (GSEA) of the Hallmark category was performed. RNAseq data are available on GEO, accession number GSE298922.

### Adhesion assay

96-well plates were coated with ECM proteins diluted in PBS for 2 hours at room temperature, then blocked with 3% filtered BSA in PBS for 15 minutes. 50,000 cells were seeded per well. At indicated timepoints, wells were washed in PBS and fixed in Crystal Violet solution (0.1% crystal violet, 20% methanol in water). Plates were incubated at 4°C overnight, rinsed in milliQ water, and incubated on shaker in the dark with 100 µL 0.1% Triton X-100 per well. Absorbance was measured at 595 nm using a SpectraMax plate reader.

### Traction-force measurements

Cells were seeded 12 days after 4OHT treatment on fibronectin-coated 30kPa polyacrylamide gels containing fluorescent nanobeads (Invitrogen). Single cells were imaged using the Nikon CSU-W1 inverted microscope, brightfield images being acquired simultaneously with nanobead fluorescence images using a ×40 objective.

Cells were then trypsinised and fluorescent images of beads were reacquired to record their position in the relaxed state. The gel deformation caused by the cells was analysed by comparing the bead positions with and without cells using a previously described Fourier transform algorithm (Butler *et al*, 2002; Trepat *et al*, 2009). The average force per unit area exerted by each cell was then calculated.

### Polyacrylamide gels for cell culture

Polyacrylamide gels of varying stiffness (Elosegui-Artola *et al*, 2016) were cast on a Bind Silane (1:14 Bind Silane solution (Sigma, M6514), 1:14 acetic glacial acid made up in 96% ethanol) treated glass surface, using a Repel Silane (Sigma, GE17-1332-01) treated coverslip to ensure a flat and even surface. Gel stiffness was adjusted by varying concentrations of 40% acrylamide and 2% bis-acrylamide in a PBS solution with 1:200 10% APS and 1:2000 TEMED. Gels were crosslinked in 5 mg/mL Sulpho-SANPAH (Sigma, 803332) in DMSO under UV for 10 minutes, and coated with 50 µg/mL fibronectin at 4°C overnight.

### Micropillars and micropatterns

For micropillars, a silicon wafer was spin-coated with SU-8 2005 photoresist at 1500 rpm to achieve ∼6 µm tall features. Soft baking was at 65°C for 1 min, then 95°C for 3 min. Photolithography was done at 120mJ/cm^2^ using an ML3 Microwriter to expose a pillar pattern of 1µm diameter, 3µm spacing, designed in Clewin. Post-exposure baking was at 65°C for 1 min, 95°C for 3 min, and 65°C for 1 min. The wafer was developed in PGMEA for 40 min, rinsed in isopropanol, dried with N_2,_ and hard baked at 200°C for 20 min. PDMS (SYLGARD 184, 10:1 base:curing agent) was applied to the patterned wafer and spin-coated under vacuum to form a thin layer. After degassing, the PDMS was cured at 110°C for 5 min, peeled off, and placed (pillars up) onto plasma-cleaned coverslips. A final bake at 110°C for 15 min was performed. Pillars were coated with 50 µg/mL fibronectin overnight.

For micropatterning, coverslips were cleaned in 1M HCl, plasma treated, and incubated with near IR labelled PLL-PEG (PLL(20)-g[3.5]-PEG(2)/Atto633). Micropatterns were generated in the PEG layer using a 185nm UV lamp (UVO 342-220, Jelight) with a custom-made quartz photomask containing the desired patterns (Compugraphics). The UV-exposed surface was coated overnight with 50 µg/mL fibronectin in NaHCO_3_ prior to cell seeding.

### OP-puromycin incorporation for protein biosynthesis

Protein biosynthesis was measured using Click-&-Go Plus OPP Protein Synthesis Assay Kit (Click Chemistry Tools, 1493). Cells were pulsed with 20 µM O-propargyl-puromycin (OPP) for 30 minutes, fixed in 4% PFA and permeabilized in 0.5% Triton X-100. After the OPP-Alexa Fluor 488 click reaction nuclei were stained with Hoechst. Coverslips were mounted in mowiol and imaged using a Zeiss Microscope AXIO Observer.D1 equipped with an AxioCam MRm camera. In ImageJ, cells were segmented using the nuclear stain for marker-controlled watershed segmentation and the mean OPP intensity per cell was measured.

## RESULTS 23-11

### The MRTFs are required for proliferation of SV40-immortalised MEFs

To investigate the role of the MRTF in cell proliferation, we established pools of SV40-immortalised mouse embryonic fibroblasts (MEFs) from animals expressing tamoxifen-inducible Cre in either wildtype (*Rosa26^Tam-Cre^*) or conditional *Mrtf*-null (*Mrtfa^-/-^*;*Mrtfb^fl/fl^*;*Rosa26^Tam-Cre^*) or conditional *Srf*-null (*Srf^/fl/fl^*;*Rosa26^Tam-Cre^*) backgrounds. All the MEF pools grew at comparable rates. To induce gene inactivation, cells were treated with 4-Hydroxytamoxifen (4OHT), and plated for analysis 7 or 10 days later (Figure 1A). Unless otherwise stated, we will hereafter refer to 4OHT-treated *Mrtfa^-/-^*;*Mrtfb^fl/fl^*;*Rosa26^Tam-Cre^*or *Srf^fl/fl^*;*Rosa26^Tam-Cre^* cells as *Mrtfab*^-/-^ and *Srf*^-/-^ MEFs.

**Figure 1:**
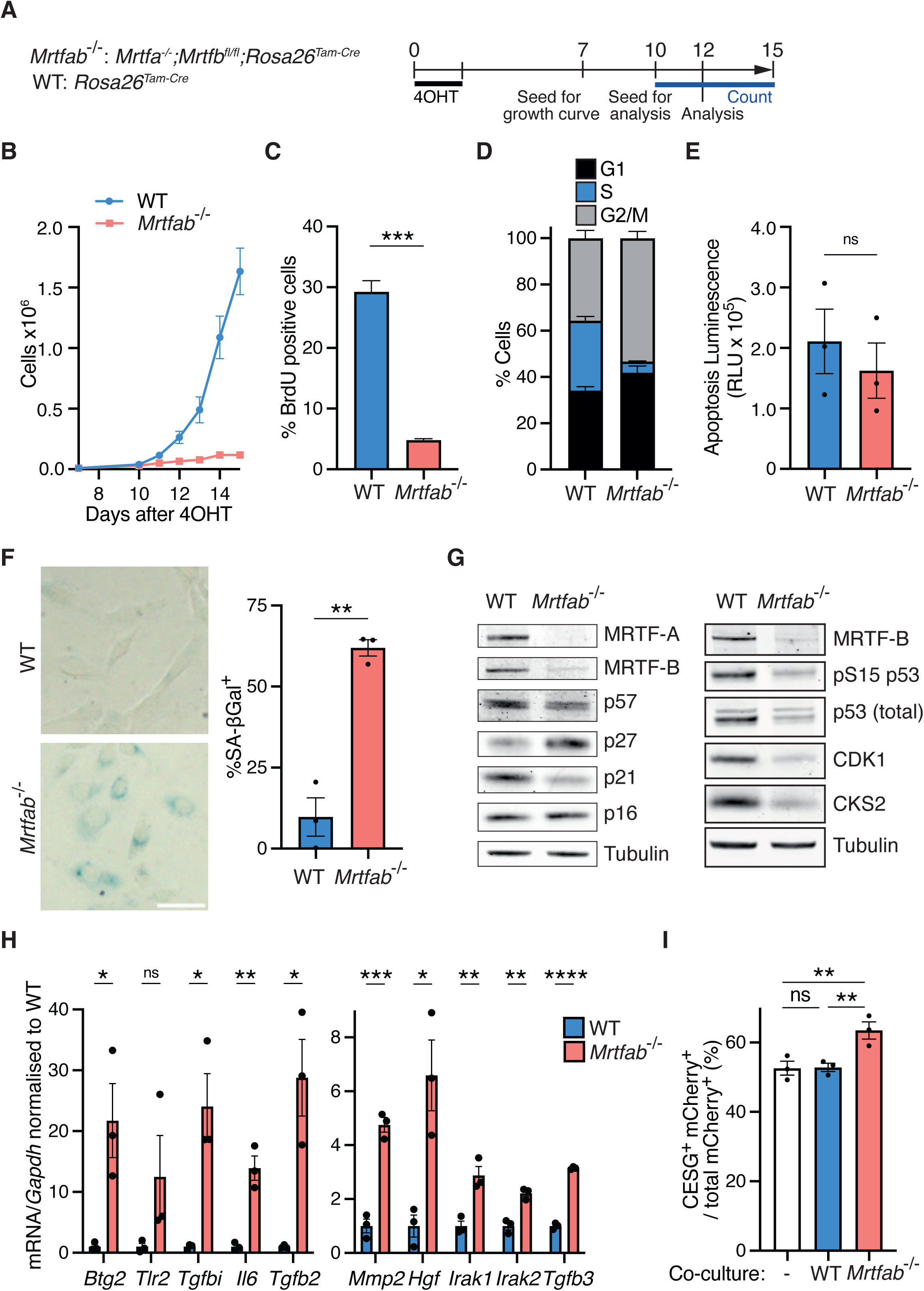
MRTF inactivation induces proliferation defect in SV40-immortalised MEFs. A) Experimental protocol. Three independent pools of SV40-immortalised MEFs were established from either *Mrtfab*^-/-^ (*Mrtfa*^-/-^;*Mrtfb*^fl/fl^;*Rosa26*^Tam-Cre^) or wildtype (*Rosa26*^Tam-Cre^) animals (3 embryos per genotype: *Mrtfab*^-/-^, pools 84, 86, 89; WT, pools 79, 80, 82). Each pool was treated with 4-hydroxytamoxifen (4OHT) for 2 days, and seeded either 7 days later for growth curves, or 10 days later for further analyses at day 12. B) Cell proliferation analysed following plating of 10,000 cells on day 7 after 4OHT treatment. For each pool, cell counts at each time point are the mean of three technical replicates. Data points are the mean cell counts of the three independent cell pools ± SEM. A representative experiment is shown. C) Quantification of BrdU-positive cells as measured by flow cytometry after a 2- hour BrdU pulse on day 12 after 4OHT. Significance, unpaired t-test. ns, not significant; *, p<0.05; **, p<0.01; ***, p<0.001; ****, p<0.0001. D) Cell cycle distribution of day 12 cells evaluated by PI/BrdU staining. Data are mean values from the three pools ±SEM. Statistical significance, Fisher’s LSD test: G1, p<0.05; S, p<0.0001; G2/M, p<0.01. E) Quantification of apoptosis in day 12 WT and *Mrtfab*^-/-^ MEFs by caspase 3/7 luminescence assay. Significance, unpaired t-test; ns. F) SA-βgal staining of day 12 WT and *Mrtfab*^-/-^ MEFs. Left, representative images of each genotype (pools 80 and 86). Scale bar = 50µm. Right, mean scores of the three pools (>50 cells scored per pool) ± SEM. Statistical analysis, unpaired t- test; p<0.01. A representative experiment is shown. G) Representative immunoblot analysis of cell cycle markers in day 12 WT (pool 80) or *Mrtfab*^-/-^ (pool 86) MEFs. For individual pool images see Figure S1A. H) RT-qPCR of mRNA levels of SASP factors in day 12 WT or *Mrtfab*^-/-^ MEFs after 4OHT. Data are mean of the three independent pools ± SEM; significance, unpaired t-test; *, p<0.05; **, p<0.01; ***, p<0.001; ****, p<0.0001. I) Primary kidney fibroblasts expressing mCherry were cultured either alone or at 100:1 with WT or *Mrtfab*^-/-^ MEFs and cultured for 5 days before staining with CellEvent Senescence Green and analysis by flow cytometry. Data are mean values for co-culture with no cells, or with each of the three independent pools of each genotype, ± SEM; statistical significance, one-way ANOVA with multiple comparisons. ns, not significant; **, p<0.01.

Untreated immortalised *Mrtfa^-/-^*;*Mrtfb^fl/fl^*;*Rosa26^Tam-Cre^*MEFs grew at similar rates to wildtype MEFs, indicating that MRTF-A is dispensable for growth. However, following 4OHT treatment all three pools of *Mrtfa^-/-^*;*Mrtfb^fl/fl^*;*Rosa26^Tam-Cre^*MEFs exhibited a severe proliferation defect compared to the wildtype MEF pools (Figure 1B). At day 12, BrdU labelling was significantly reduced, while proportions of both G1 and G2/M cells increased, the proportion of apoptotic cells remaining unchanged (Figure 1C-E). The *Mrtfab^-/-^* cells also had elevated levels of neutral β-galactosidase (SA-βgal) activity, conventionally regarded as a senescence marker (Dimri *et al*, 1995)(Figure 1F). Replicative senescence is generally associated with upregulation of the cyclin-CDK inhibitors (CKIs) CDKN1A/p21 and CDKN2A/p16, which inhibit the pro-proliferative Cdk4/6-Rb-E2F pathway (Campisi, 2013). Immunoblot analysis of CKI and CDK expression in *Mrtfab^-/-^*MEFs revealed upregulation of p27, while p21 and cyclin D1 were downregulated, as were p53, CDK1 and CKS2, while p16 was unchanged (Figure 1G, Figure S1A). Similar results were obtained for each *Mrtfab^-/-^* MEF pool, and without 4OHT treatment, *Mrtfa^-/-^*;*Mrtfb^fl/fl^*MEFs displayed a similar profile to wildtype cells (Figure S1A, S1B). Previous studies have demonstrated that MRTF-SRF signalling suppresses ERK activity and oncogene-induced senescence in *DLC1*-deleted cancer cells (Hampl *et al*., 2013), but in contrast, *Mrtfab^-/-^* MEFs exhibited reduced ERK activity (Figure S1C).

*Mrtfab^-/-^* MEFs exhibited elevated expression of senescence-associated secretory phenotype (SASP) factors (Acosta *et al*., 2013), including secreted proteins (*Il6*, *Mmp2*, *Hgf*), cellular receptors (*Tgfb2*, *Tgfb3*, *Tgfbi*, *Tlr2*) and downstream effectors (*Btg2*, *Irak1*, *Irak2*) (Figure 1H, Figure S1A). The SASP can act in a paracrine fashion to induce senescence (Acosta *et al*., 2013). We cultured *Mrtfab^-/-^*or WT cells with mCherry-expressing primary kidney fibroblasts for 5 days, and assessed senescence by staining with CellEvent Senescence Green (CESG). Co-culture with *Mrtfab^-/-^* MEFs, but not wildtype MEFs, increased the proportion of senescent CESG+ primary kidney fibroblasts, indicating that *Mrtfab^-/-^* cells can act in a paracrine manner to influence fibroblast proliferation (Figure 1I). Since the MRTFs act in partnership with SRF, we also analysed *Srf*^-/-^ MEFs. These cells behaved similarly to *Mrtfab*^-/-^ MEFs, exhibiting defective proliferation, SA-βGal expression, altered CKI expression, and upregulation of SASP markers (Figure S1D-H).

### MRTFs are required for proliferation of primary fibroblasts and epithelial cells

To investigate whether the proliferative defect phenotype observed in immortalised *Mrtfab^-/-^* MEFs was also seen in primary fibroblast cells, we isolated lung and kidney fibroblasts from *Mrtfa^-/-^;Mrtfb^fl/fl^;Rosa26^Tam-Cre^* or wildtype *Rosa26^Tam- Cre^* animals and treated them with 4OHT to induce MRTF inactivation (Figure 2A). *Mrtfab^-/-^* lung and kidney fibroblasts both exhibited a clear proliferative defect compared with their wildtype counterparts (Figure 2B), and also stained positive for SA-βGal (Figure 2C). The lung fibroblasts behaved similarly to the immortalised MEF pools, with the p27 CKI and the SASP regulator TLR2 upregulated (Figure 2D), and displayed increased levels of *Hgf*, *Mmp3*, and *Il6* mRNAs (Figure 2E). In contrast, kidney fibroblasts displayed upregulation of markers conventionally associated with oncogene-induced senescence, including phospho-p53 and the CKIs p21 and p16, and upregulated the SASP components *Tlr2*, *Btg2*, *Irak1*, *Mmp9*, *Il6*, and *Il1a* (Figure 2D, 2E). Thus, inactivation of the MRTFs induces a proliferation defect with senescence-like characteristics in primary kidney and lung primary fibroblasts.

**Figure 2:**
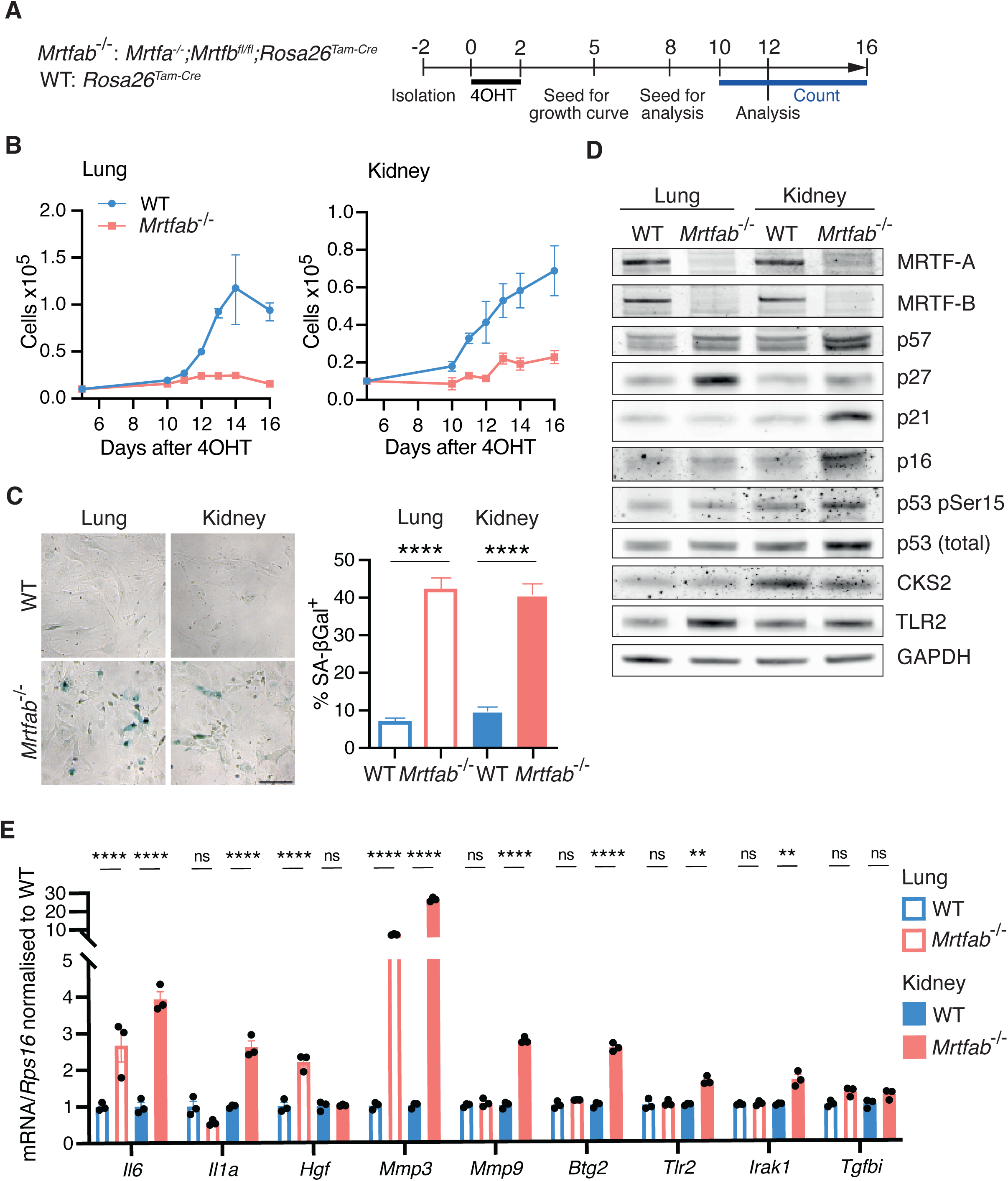
Defective proliferation of MRTF-null primary tissue-derived fibroblasts. A) Experimental protocol. Fibroblast pools were isolated from lung and kidney tissue of single WT or *Mrtfa^-/-^*;*Mrtfb^fl/fl^* mice and treated two days later with 4OHT, before seeding for analysis as indicated. B) 10,000 cells seeded on day 5 after 4OHT and counted from day 10 to 16. N=3 mice. Data points are the mean cell counts of three independent isolates of each genotype ± SEM. C) SA-βgal staining on day 12 after 4OHT. Scale bar = 50µm. Right, mean scores of five isolates per genotype (>100 cells scored each isolate) ± SEM with one-way ANOVA with multiple comparisons for statistical analysis; p<0.0001. D) Immunoblotting of lung and kidney fibroblasts on day 12 after tamoxifen. Representative of three independent experiments. E) RT-qPCR of mRNA levels of SASP factors in lung and kidney fibroblasts on day 12 after 4OHT. Data is from a single experiment with 3 technical replicates, ± SEM, with two-way ANOVA with multiple comparisons for statistical analysis: ns, not significant; **, p<0.01; ****, p<0.0001.

We also examined the role of MRTF-SRF signalling in proliferation of epithelial cells. Primary tracheal epithelial cells (MTECs) were isolated from *Mrtfa^-/-^; Mrtfb^fl/fl^;Rosa26^Tam-Cre^* or wildtype *Rosa26^Tam-Cre^* animals using an established protocol (Crotta *et al*., 2023; You *et al*., 2002), treated with 4OHT, and plated on transwell plastic inserts (Figure S2A). Upon culture under non-differentiating conditions, both wildtype and *Mrtfab^-/-^*cells formed a non-ion permeable barrier, as judged by trans-epithelial resistance, although that formed by *Mrtfab^-/-^* cells appeared somewhat weaker (Figure S2B). Strikingly, however, although wildtype cells underwent several divisions after this point, *Mrtfab^-/-^*cells greatly reduced proliferation (Figure S2C), accumulating predominantly in G2/M with increased SA-βGal activity and SASP marker expression (Figure S2D-S2F). This was accompanied by elevated p21 and decreased p27 expression, and CDK1 and CKS2 levels were reduced, as seen in the fibroblasts (Figure S2G). MRTF-SRF signalling is thus required for effective proliferation of both primary fibroblasts and epithelial cells.

### The *Mrtfab^-/-^* proliferation defect can be reversed by MRTF-A re-expression

Proliferative defects, accompanied by elevated SA-βGal activity and SASP marker expression, are phenotypes generally associated with cell senescence, which is conventionally regarded as irreversible. We therefore next tested whether the phenotypes of *Mrtfab^-/-^*MEFs could be reversed by MRTF-A re-expression. Clonal cell lines expressing doxycycline (Dox)-inducible HA-tagged MRTF-A together with an IRES-GFP expression marker, were derived from *Mrtfa^-/-^; Mrtfb^fl/fl^;Rosa^Tam-Cre^* clone 86 MEFs cells by lentiviral infection, and analysed with and without Dox and/or 4OHT treatment (hereafter *Mrtfa^-/-^*;*Mrtfb^fl/fl^*DoxMRTF-A cells, Figure 3A). 4OHT treatment, which inactivates endogenous *Mrtfb*, impaired *Mrtfa^-/-^*; *Mrtfb^fl/fl^* DoxMRTF-A proliferation, as expected, but this did not occur when MRTF-A expression was induced by Dox treatment 7 days later (Figure 3A). MRTF-A re-expression partially suppressed the increases in TLR2 and p27, and decreases in CDK1 and CKS2 expression (Figure 3B). It also reduced SA-βGal activity (Figure 3C) and transcription of SASP markers (Figure S3A).

**Figure 3:**
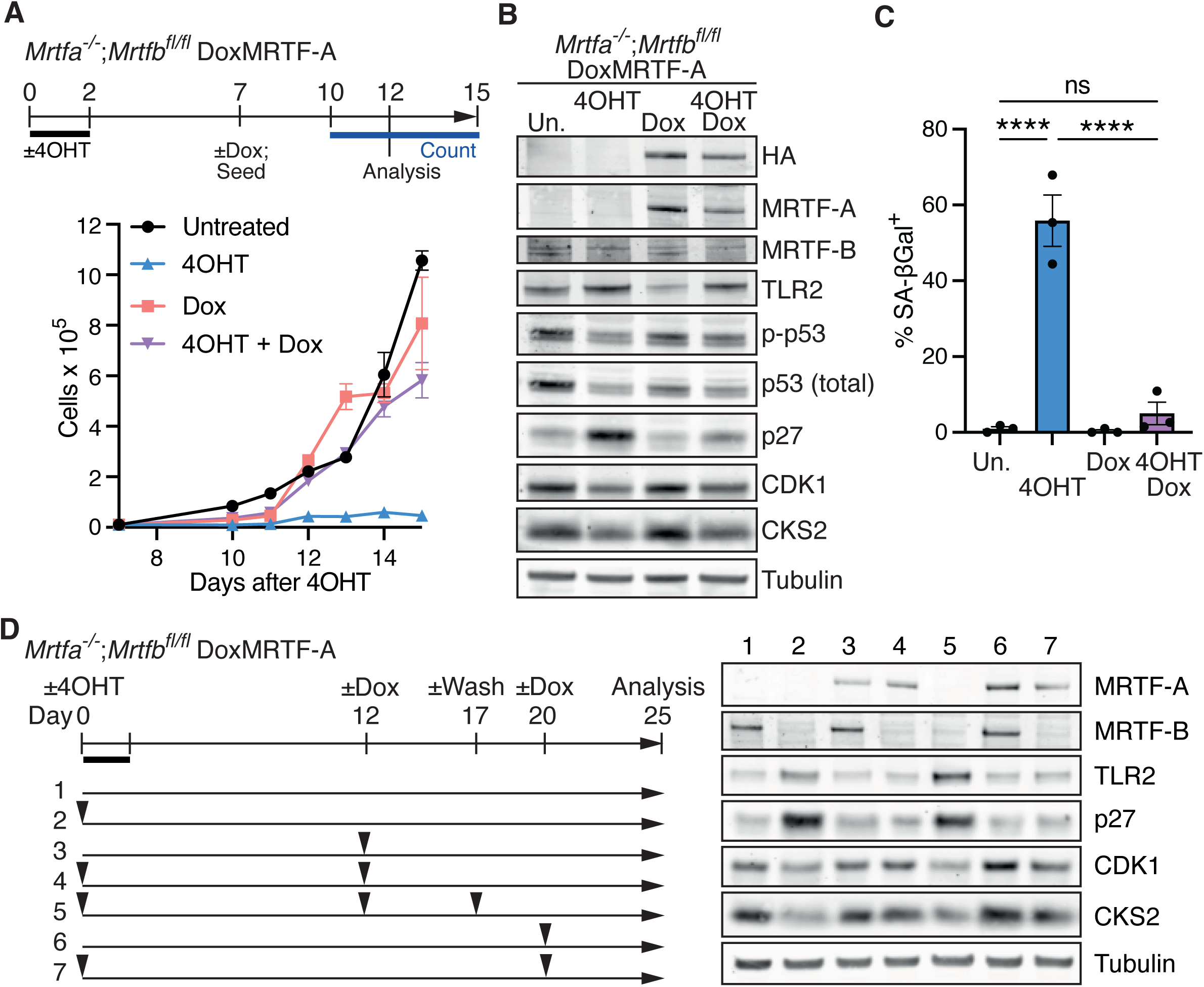
Proliferation of *Mrtfab*^-/-^ MEFs is reversed by MRTF-A re-expression. A) Top, experimental protocol. Three independent *Mrtfa*^-/-^*Mrtfb*^fl/fl^DoxMRTF-A clonal cell lines, derived from Pool 86 *Mrtfa*^-/-^*Mrtfb*^fl/fl^*Rosa26*^Tam-Cre^ MEFs, were treated or not with 4OHT and/or doxycycline 7 days later. Bottom, cell proliferation following seeding of 10,000 cells of each line in triplicate at day 7. Cell counts were determined for cell line 1F2 as the mean of three technical replicates ± SEM. Similar results were obtained with lines 1F5 and 1B2. B) Cell cycle markers in *Mrtfa^-/-^*;*Mrtfb^fl/fl^* DoxMRTF-A cell line 1F2 treated as in (A) at day 12 were analysed by immunoblotting. Similar results were obtained with lines 1F5 and 1B2. C) SA-βgal staining of day 12 *Mrtfa^-/-^Mrtfb^fl/fl^* DoxMRTF-A cells treated as in (A). at day 12. Data are mean scores of the three lines (>50 cells scored per pool) ± SEM. Statistical analysis, one-way ANOVA corrected for multiple comparisons; ns, not significant; ****, p<0.0001. D) Extended MRTF-A activation and shutoff. Left, experimental protocol. *Mrtfa^-/-^ Mrtfb^fl/fl^* DoxMRTF-A-HA cells were treated or not with 4OHT and/or doxycycline and/or doxycycline washout at the indicated times. Right, cell cycle marker expression in line 1F2 at day 25. Similar results were obtained with line 1F5.

We explored the effect of MRTF-A re-expression and shut-off at different times after MRTF inactivation (Figure 3D). Re-expression of MRTF-A at 12 or even 20 days after MRTF inactivation restored p27, TLR2, CDK1 and CKS2 levels to those seen in untreated cells (Figure 3D, samples 3, 6). Moreover, these effects were themselves reversible: the changes induced by MRTF-A re-expression at 12 days could be reversed by Dox washout at day 17 (Figure 3D, sample 5). MRTF-A re-expression also suppressed SA-βGal activity and SASP marker expression (Figure S3B, S3C). Taken together, these results show that the proliferation defect is reversible, and that MRTF-A functions redundantly with MRTF-B to promote cell proliferation.

### Serum-deprivation induces a similar proliferative defect to MRTF inactivation

The reversibility of the proliferation defect associated with MRTF inactivation led us to investigate its relationship with the proliferative arrest induced by serum deprivation. Wildtype or *Mrtfa^-/-^*;*Mrtfb^fl/fl^*cells were treated with 4OHT and then cultured in 0.3% or 10% serum prior to cell proliferation analysis (Figure 4A). Upon culture in 0.3% serum, wildtype cell proliferation became comparable to that of *Mrtfab^-/-^* cells cultured in 10% serum, while *Mrtfab^-/-^* proliferation became virtually undetectable (Figure 4B). After five days of serum deprivation, WT MEFs exhibited substantially increased expression of p27, and decreased CDK1, CKS2 and p53 phospho-S15 to levels similar to those seen in *Mrtfab^-/-^* MEFs maintained in 10% serum (Figure 4C). Similar but less pronounced changes were seen after 2 days of serum deprivation (Figure S4A). Serum deprivation induced SA-βGal activity in WT cells, and significantly increased it in *Mrtfab^-/-^* cells (Figure 4D). Moreover, SASP factor expression was also increased in serum-deprived WT cells, as assessed by qRT-PCR (Figure S4B). Both quiescence and senescence are associated with global decreases in protein translation (reviewed by Marescal & Cheeseman, 2020; Payea *et al*, 2021). We found that *Mrtfab^-/-^* cells cultured in 10% serum displayed a diminished protein synthesis rate compared to WT MEFs as assessed by O-propargyl-puromycin (OPP) incorporation; in both cases, culture in 0.3% reduced global protein synthesis rate further, to a similar basal level (Figure 4E).

**Figure 4:**
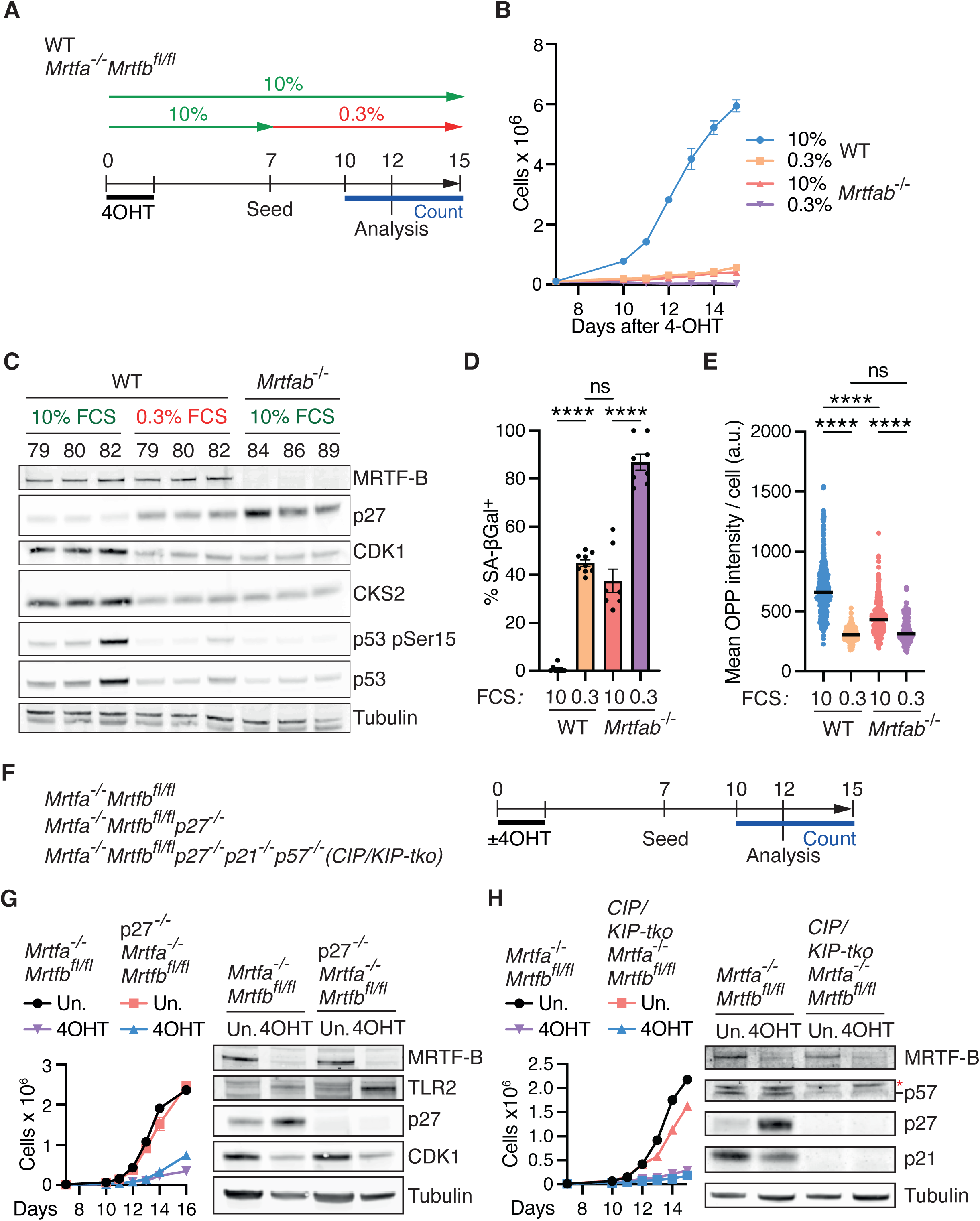
The MRTF-null phenotype exhibits features of quiescence. A) Experimental protocol. The three pools of WT and *Mrtfab*^-/-^ MEFs were treated with 4OHT and cultured in 10% FCS, with medium replaced or changed to 0.3% FCS on day 7. B) Proliferation of cells treated as in (A). 10,000 cells were seeded on day 7. Data are mean ± SEM of the 3 pools for each genotype. C) The three pools of WT and *Mrtfab*^-/-^ MEFs were treated as in (A) and cell cycle markers analysed by immunoblot at day 12. D) WT (pool 80) and *Mrtfab*^-/-^ (pool 86) MEFs were treated as in (A) and SA-βGal staining analysed at day 12. Data are mean ± SEM, n=7 to 9 replicates per line, with one-way ANOVA for statistical analysis. ns, not significant; *, p<0.05; **, p<0.01; ***, p<0.001; ****, p<0.0001. E) WT (pool 80) and *Mrtfab*^-/-^ (pool 86) MEFs were treated as in (A) and protein biosynthesis measured by O-propargyl-puromycin incorporation at day 12. Data are values for individual cells (>170 cells per condition/genotype), with mean indicated. One-way ANOVA was used for statistical analysis. ns, not significant; ****, p<0.0001. F) p27-null and CIP/KIP-null *Mrtfa*^-/-^*Mrtfb*^fl/fl^ MEFs were generated from *Mrtfa*^-/-^;*Mrtfb*^fl/fl^ pool 86 MEFs by CRISPRCas9 mutagenesis. Following treatment or not with 4OHT, 10,000 cells were seeded on day 7, with immunoblot analysis at day 12. G) Growth curve and cell cycle markers for p27^-/-^*Mrtfa*^-/-^*Mrtfb*^fl/fl^ MEFs treated as in (F). Data for clone 3A2 are shown. Similar results were obtained with clones 1F3 and 3B10. H) Growth curve and cell cycle markers for clone p27^-/-^p21^-/-^p57^-/-^*Mrtfa*^-/-^*Mrtfb*^fl/fl^ MEFs treated as in (F). Data for clone 3.1D10 are shown. Similar results were obtained with clones 3.1C11 and 3.1G9.

Previous studies have demonstrated that p27 inactivation does not affect the density at which primary MEFs arrest in culture, or their sensitivity to serum deprivation (Nakayama *et al*, 1996). We therefore tested whether inactivation of p27 could relieve the proliferation defect in *Mrtfab^-/-^* MEFs. We used CRISPR-Cas9 to inactivate p27 in *Mrtfa^-/-^*;*Mrtfb^fl/fl^* cells, evaluating proliferation and marker expression with and without 4OHT treatment. Inactivation of p27 did not relieve the defective proliferation seen upon MRTF inactivation (Figure 4F,G), and as seen in *Mrtfab^-/-^* MEFs,p27*^-/-^Mrtfab^-/-^*MEFs exhibited reduced CDK1 expression, elevated SA-βGal and SASP factor expression (Figure 4G, S4C,D). The CDK inhibitor p21/*Cdkn1a* is also implicated in MRTF-SRF controlled proliferation, at least in the settings described above, and p27 is at least to some extent functionally redundant with p57/*Cdkn1c* in cell cycle control (Matsumoto *et al*, 2011; Susaki *et al*, 2009). Nevertheless, inactivation of all three CDK inhibitors in *Mrtfa^-/-^Mrtfb^fl/fl^* cells also failed to relieve the proliferation defect resulting from MRTF inactivation (Figure 4H). Conversely, overexpression of CDK1, CKS2, or myoferlin using the doxcycline- inducible retroviral transduction strategy used for MRTF-A re-expression failed to rescue proliferation of *Mrtfab^-/-^* MEFs (Figure S4E).

Taken together with the results in the preceding section, these data show while the proliferative defect in *Mrtfab^-/-^* MEFs is substantially similar to that induced by serum deprivation, and that the altered expression of cell cycle regulators associated with it are likely to be its consequence rather than its cause.

### Cytoskeletal and cell cycle genes are downregulated in *Mrtfab^-/-^* MEFs

MRTF-SRF signalling plays a central role in regulation of cytoskeletal gene expression (Esnault *et al*., 2014; Olson & Nordheim, 2010; Schratt *et al*., 2002). We used RNAseq to analyse differentially expressed mRNAs in immortalised wildtype and *Mrtfa^-/-^*;*Mrtfb^fl/fl^*MEFs at 0, 2, 4, and 12 days following treatment with 4OHT (referred to as T0, T2, T4, T12 cells, Figure S5A). Differential gene expression analysis showed that before 4OHT treatment, *Mrtfa^-/-^*;*Mrtfb^fl/fl^*MEFs, which express MRTF-B alone, exhibited a very similar gene expression profile to wildtype cells (Figure 5A, Table 1; 119 genes down, 70 up at log2 foldchange = 1, Padj <0.05). These genes did not exhibit significant enrichment in GSEA analysis (Figure S5B).

**Figure 5:**
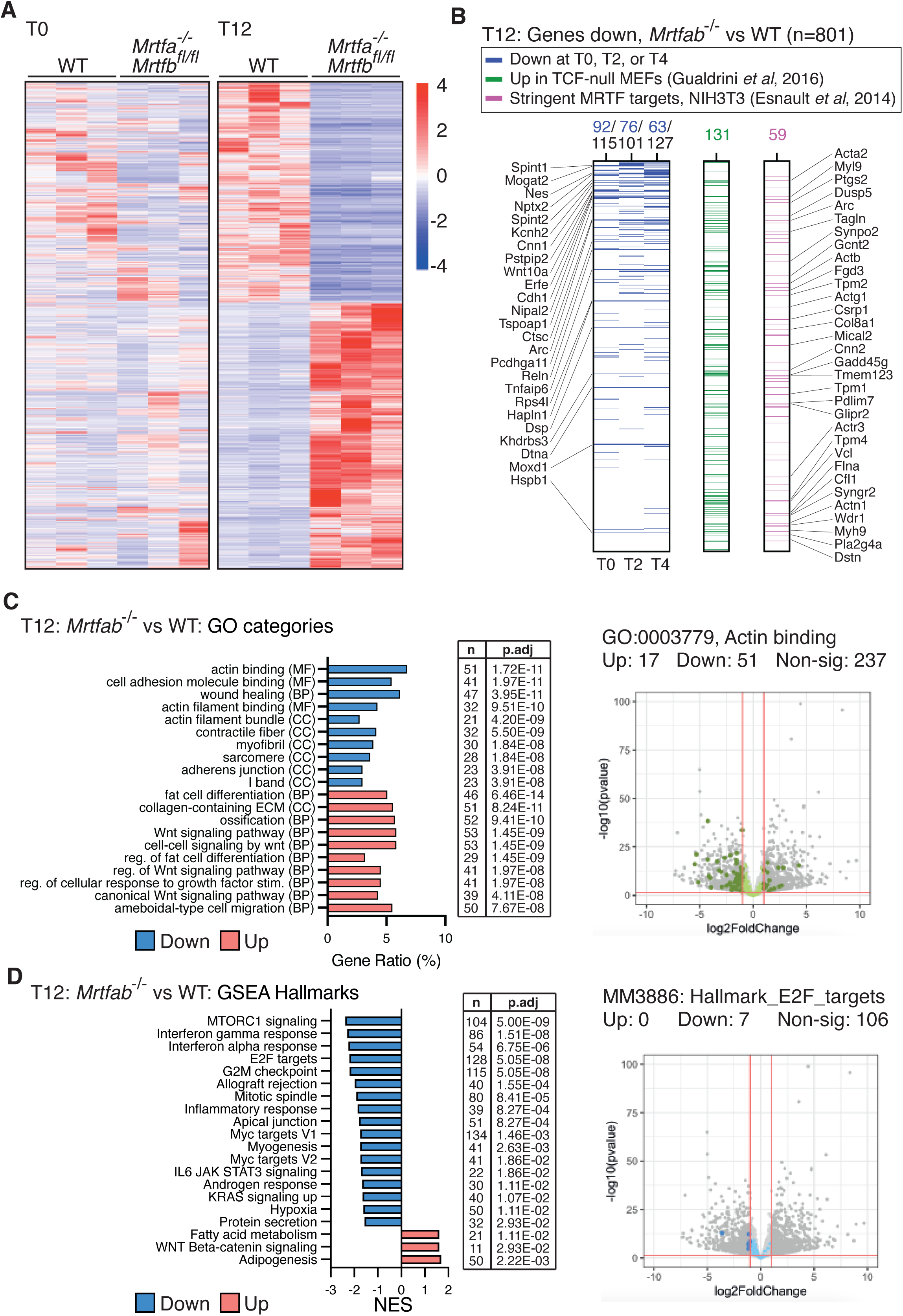
Downregulation of cytoskeletal and cell cycle genes in *Mrtfab^-/-^* MEFs. A) WT MEF pools 79, 80, 82; and *Mrtfa*^-/-^*Mrtfb^fl/fl^* MEF pools 84, 86, 89 were treated or not with 4OHT and cultured for 0, 2, 4, or 12 days before RNAseq analysis (T0, T2, T4, T12 samples, Table 1; see Figure S5A). Heatmap shows genes differentially expressed at T12 (>2-fold change and Padj < 0.05 in the comparison between WT and *Mrtfab^-/-^* MEFs at T12). z-scores of genes for T0 and T12 cells are ordered using unsupervised clustering. RNAseq data are deposited on GEO, accession number GSE298922. B) Relationships between the 801 genes with gene symbols that map to an ensembl ID and are significantly downregulated (>2-fold change, Padj <0.05) in *Mrtfab^-/-^* MEFs on day 12. The genes are ranked by fold-change upon MRTF inactivation. Left (blue), genes downregulated in *Mrtfab^-/-^* MEFs at other timepoints, with those common to all timepoints listed; centre (green), identity with genes upregulated in MEFs upon inactivation of the TCFs (Gualdrini *et al*., 2016); right (magenta), identity with the stringently defined MRTF target genes in NIH3T3 cells (Esnault *et al*., 2014); list shows genes in common with both datasets. C) GO category analysis of differentially regulated genes in *Mrtfab^-/-^* MEFs on day 12. Bar plots, top 10 downregulated (blue) or upregulated (red) GO terms from any category (BP: Biological Process, MF: Molecular Function, CC: Cellular Component) ranked according to Padj. Table, number of genes changing, and the Benjamini-Hochberg adjusted p-value for each gene set. The specimen GO term GO:0003779 Actin Binding is shown at right. D) GSEA Hallmark analysis of differentially regulated genes in *Mrtfab^-/-^* MEFs on day 12. Bar plots, top 20 GSEA hallmark gene sets, showing the normalised enrichment score (blue, downregulated; red, upregulated). Table, number of genes changing, and the Benjamini-Hochberg adjusted p-value for each gene set. The specimen GSEA Hallmark MM3886: HALLMARK_E2F_TARGETS is shown at right.

Although little change was observed at early times after 4OHT treatment by day 12 the resulting *Mrtfab^-/-^* MEFs exhibited substantial changes in gene expression (Figure 5A, Figure S5C) (801 genes down, 1065 up; see Table 1). Significantly down-regulated genes included 59 out of the 683 genes identified as MRTF-SRF targets in serum-stimulated NIH3T3 cells (p= 2.2e-07, Esnault *et al*., 2014), and 131 of 328 putative MRTF target genes whose expression in MEFs is repressed by the TCF cofactors (p= 1.9e-13, Gualdrini *et al*, 2016); a substantial number of genes (32/801) were found in all three datasets (Figure 5B). GO analysis revealed downregulation of members of genes involved in the structure and regulation of the actin cytoskeleton and its functions, such as adhesion and migration (Figure 5D, Table 2). In addition, T12 *Mrtfab^-/-^*cells downregulated genes associated with GSEA Hallmarks involved in cell proliferation, cell cycle progression, and growth (Figure 5E, Table 2).

Previous studies that have suggested that MRTF-SRF signalling suppresses cell state plasticity (Hu *et al*, 2019; Zhang *et al*, 2023). Consistent with this, T12 *Mrtfab^-/-^* cells exhibited upregulation of gene sets related to adipogenesis and Wnt signalling (Figure 5D, E) and upregulation of the adipogenic markers in T12 *Mrtfab^-/-^* cells was confirmed using qRT-PCR (Figure S5D). Adipocytes could not be detected in T12 *Mrtfab^-/-^* cells by Oil Red O staining, however, although both wildtype and T12 *Mrtfab^-/-^* cells could differentiate into fat cells under adipogenic culture conditions (Figure S5E). Thus, T12 *Mrtfab^-/-^* MEFs appear to have relaxed cell state plasticity (see Discussion).

### Altered morphology and cytoskeletal dysfunction in *Mrtfab^-/-^* MEFs

We next investigated the relationship between defective cytoskeletal gene expression in *Mrtfab^-/-^* MEFs and proliferation. Consistent with studies in other systems (Alberti *et al*., 2005; Medjkane *et al*., 2009; Schratt *et al*., 2002). *Mrtfab^-/-^* MEFs exhibited a rounded morphology and lacked well-defined stress fibers and focal adhesions, which were replaced by cortical paxillin-rich puncta (Figure 6A). F-actin levels, as assessed by phalloidin staining, were significantly reduced (Figure 6B), and total actin levels were substantially decreased, as was the F/G-actin ratio (Figure 6C). MRTF-A re-expression in *Mrtfab^-/-^* MEFs was sufficient to restore normal cell morphology and F-actin distribution (Figure S6A). Phasefocus livecyte microscopy revealed that *Mrtfab^-/-^* MEFs occupy a reduced cell area, with increased sphericity and cell thickness, but their volume was unchanged (Figure 6D). Upon plating, *Mrtfab^-/-^* MEFs, spread more slowly than wildtype cells, and more isotropically (Figure S6B, S6C). They were defective in adhesion to the integrin ligands fibronectin or vitronectin, but not to Poly-L-Lysine (PLL) which allows integrin-independent cell adhesion, consistent with their lack of focal adhesions (Figure S6D). *Mrtfab^-/-^*MEFs also exhibited defective motility, with significantly reduced instantaneous velocity compared to wildtype cells and little net displacement over the 96h monitoring period (Figure 6E; Videos 1,2). MRTF-A re-expression in *Mrtfab^-/-^* DoxMRTF-A cells increased cell velocity and net displacement, and decreased cell sphericity (Figure S6E,F; Videos 3-5). Taken together, these data show that MRTF-A and MRTF-B function redundantly in MEFs to control actin cytoskeletal dynamics, adhesive and motile behaviour.

**Figure 6:**
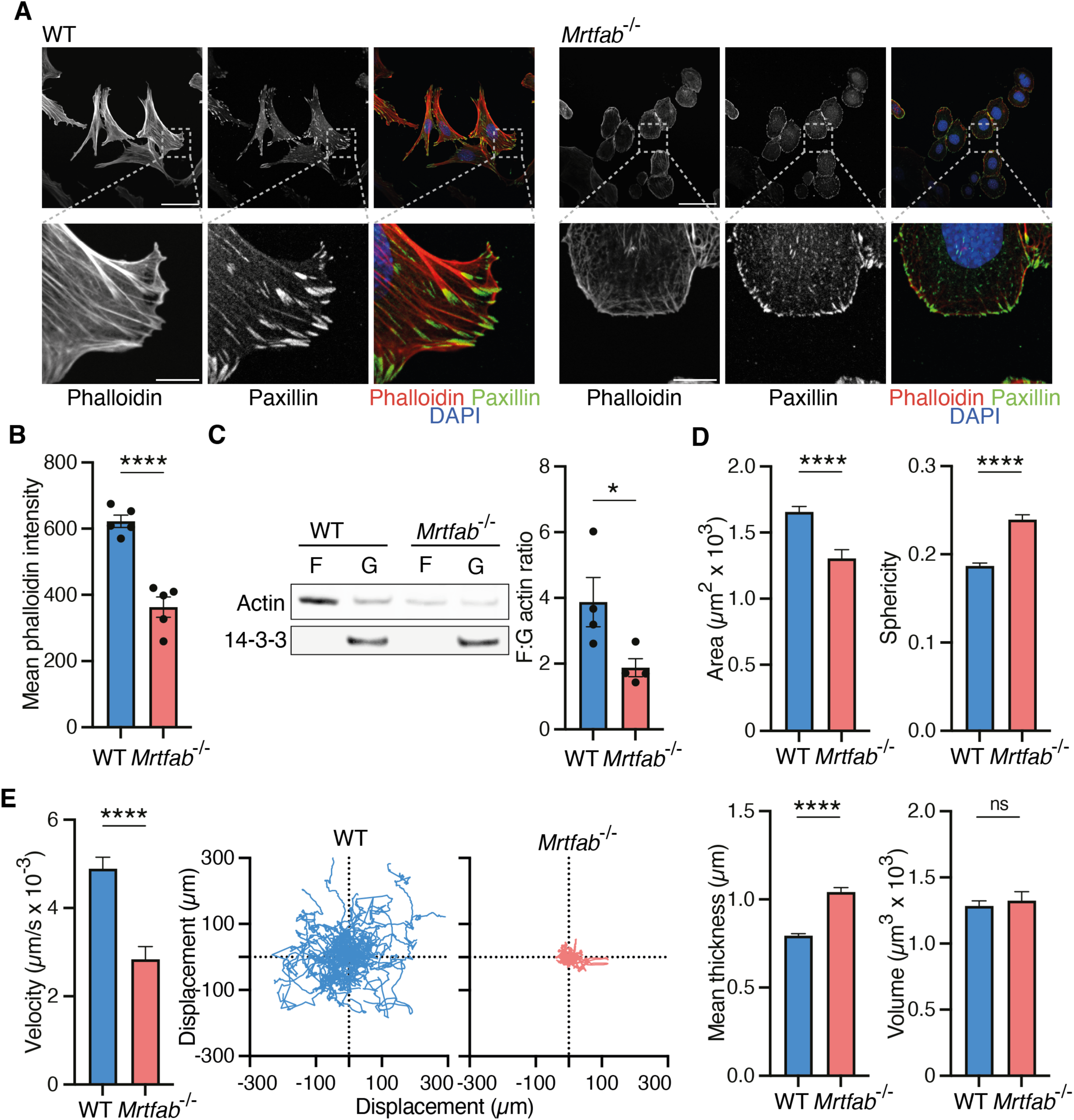
Cytoskeletal defects in *Mrtfab*^-/-^ MEF. WT or *Mrtfab*^-/-^ MEFs (pools 80 and 86) were analysed at day 12 after 4OHT. Similar results were obtained with the other MEF pools. A) Immunofluorescence staining of paxillin (focal adhesions), phalloidin (F-actin) and DAPI (DNA). Scale bar = 50µm (regular images) and 10µm (zoomed in images). Representative of 5 independent experiments. B) Mean phalloidin staining intensity per cell of cells cultured and stained as i (A). Data are mean ± SEM of 5 independent experiments. Statistical analysis, unpaired t-test; p<0.0001. C) Left, F-actin pelleting assay. 14-3-3, soluble protein control. Representative of 4 independent experiments. Right, F/G ratio; data are mean ± SEM, n = 4; statistical analysis, unpaired t-test; p<0.05. D) Morphological analysis of cell area, sphericity, thickness and volume by Phasefocus Livecyte Microscopy. Measurements are from the first frame of imaging sequence, >100 cells per sample. Data are mean from three technical replicates ± SEM. Statistical analysis, unpaired t-test; ns, not significant; ****, p<0.0001. E) Analysis of cell motility of cells in (D) by Phasefocus Livecyte Microscopy. Cells were imaged over 96 hours at 2 images/h. 50 trajectories were monitored until cell division, exit of frame of view, or end of video. Left, instantaneous velocity, plotting measurements from frame n to n+1 throughout the whole imaging sequence. >100 cells per sample with data compiled from three technical replicates. Data are mean ± SEM. Statistical analysis, with unpaired t-test; p<0.0001. Right, displacement plots of cells in a representative experiment.

### *Mrtfab^-/-^* MEF proliferation remains responsive to mechanical cues

Previous studies have shown that proliferation of adherent cells is dependent on both adhesion and matrix compliance, and cell tension and spreading (Chen *et al*, 1997; Huang *et al*, 1998; Klein *et al*, 2009; Mammoto, 2004 #102, reviewed by Mammoto & Ingber, 2009). The multiple cytoskeletal defects seen in *Mrtfab^-/-^* MEFs raises the possibility that their defective proliferation might result from inability to respond to such mechanical environmental cues. First, we examined the effect of matrix complicity. Wildtype and *Mrtfab^-/-^* MEFs were plated on substrates of increasing rigidity, and proliferation measured by EdU incorporation. Proliferation of wildtype cells was substantially reduced on soft substrates, as expected, to a rate comparable to that of *Mrtfab^-/-^*cells on hard substrates; however, *Mrtfab^-/-^* cell proliferation was further reduced on soft substrates (Figure 7A). Next, we examined the effects of cell spreading. Proliferation of wildtype cells was reduced upon plating on patterns which limit cell-substrate contact area (Figure 7B) but was unaffected when cells were grown on pillars, which restrict the total adhesion area but allow cells to spread normally (Figure 7C). Proliferation of *Mrtfab^-/-^* cells was similar to that seen for wildtype cells under conditions of limiting adhesion area; however, they remained sensitive to confinement, indicating that other pathways might also contribute (see Discussion).

**Figure 7:**
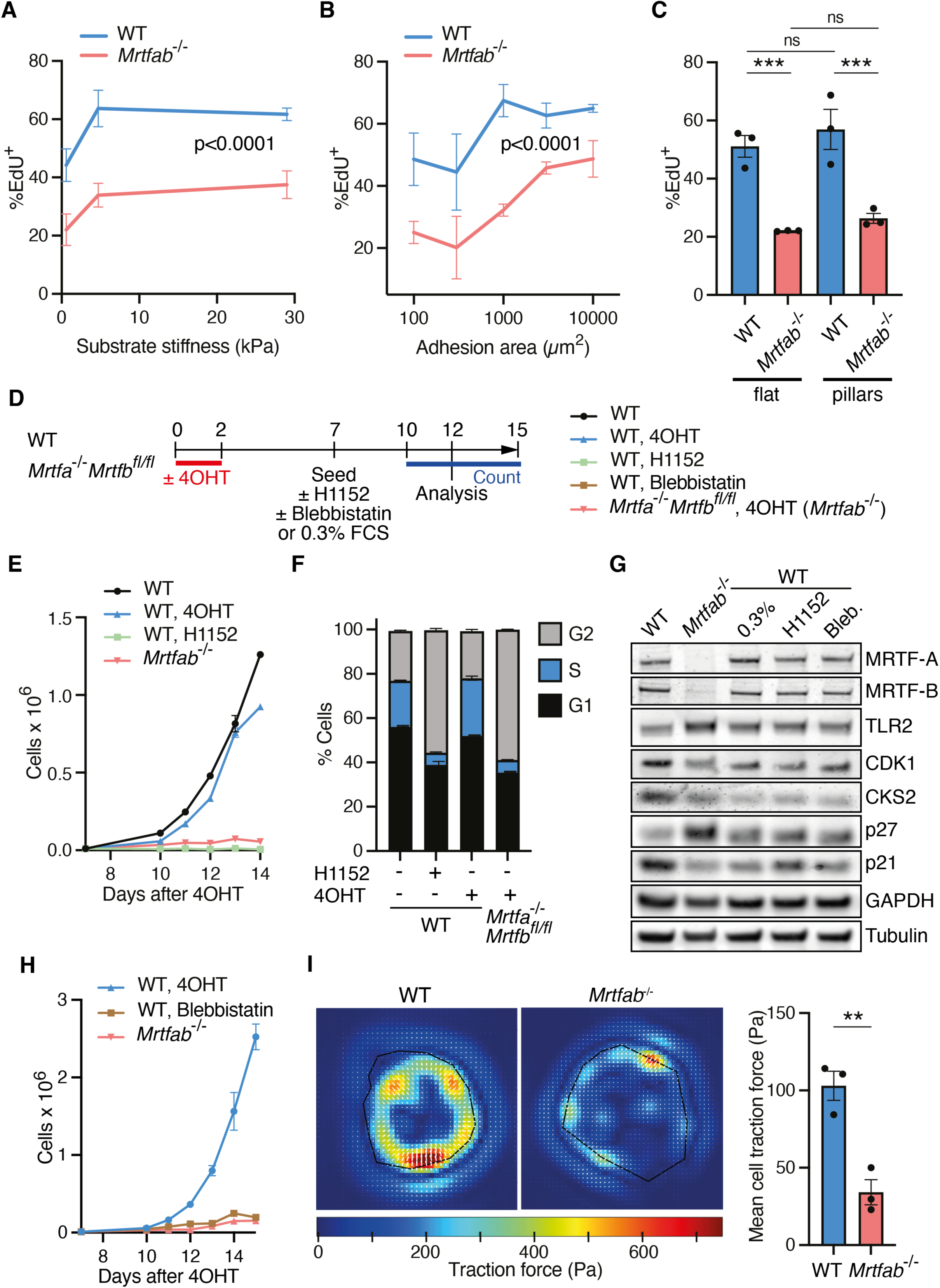
Inhibition of contractility in wildtype cells induces a phenotype similar to MRTF inactivation. A) Cells from the three WT or *Mrtfab*^-/-^ MEF pools were plated on polyacrylamide substrates of different stiffnesses at day 10 following 4OHT treatment, cultured for 2 days and cell cycle status assessed by EdU incorporation during a 6h period on day 12. Data are mean of >100 cells per sample, ± SEM. Statistical analysis by two-way ANOVA reports significance of the main effect (WT vs. *Mrtfab*^-/-^). B) Cells from each WT or *Mrtfab*^-/-^ MEF pool were plated on islands of different areas at day 10 following 4OHT treatment. Area per cell is ∼1500 μm^2^ (Figure 4D). Cell cycle status at day 12 was analysed and presented as in (A). C) Cells from each WT or *Mrtfab*^-/-^ MEF pool were plated at day 10 following 4OHT treatment either on flat surfaces or on etched patterned substrates (1µm in diameter and 3 µm between features) in which only ∼5% of a given area was available for adhesion. EdU incorporation at day 12 was analysed and presented as in (A). Statistical analysis used one-way ANOVA with multiple comparisons using Fisher’s LSD test: ns, not significant; ***, p<0.001. D) Schematic of protocol for panels E-H. *Mrtfab*^-/-^ (*Mrtfa*^-/-^*Mrtfb*^fl/fl^*Rosa26*^Tam-Cre^) pool 86 or WT (*Rosa26*^Tam-Cre^) pool 80 cells were treated or not with 4OHT, H1152 and blebbistatin as indicated before analysis. E) Growth curves of pools 80 and 86 cells treated as in (D). 10,000 cells were seeded on day 7 and counted from day 10 to 15. Data are mean of three technical replicates ± SEM, and are representative of three independent biological replicates. F) Quantification of BrdU-positive cells as measured by flow cytometry after a 2- hour BrdU pulse on day 12 after 4OHT. Data are mean of the three independent pools ± SEM; significance, unpaired t-test G) Representative immunoblot analysis of cell cycle markers in day 12 WT (pool 80) or *Mrtfab*^-/-^ (pool 86) MEFs treated as indicated. H) Growth curves of WT (pool 80) treated with blebbistatin compared with *Mrtfab*^-/-^ cells (pool 86) as in (D). 10,000 cells were seeded on day 7 and counted from day 10 to 15. Data are mean of 3 technical replicates ± SEM. I) The three pools of WT or *Mrtfab*^-/-^ MEFs were plated on 30 kPa gels and analysed by traction force microscopy on day 12. Traction force was determined for 15-20 cells for each pool. Images of deduced traction force for representative cells from pools 80 and 86. Data are mean traction force for each pool ± SEM, n = 3 independent experiments. Statistical analysis, unpaired t-test; p <0.01.

### Defective *Mrtfab^-/-^* MEF proliferation reflects impaired actomyosin contractility

Given the role of MRTF-SRF signalling in regulation of cytoskeletal dynamics, we next investigated how direct interference with cytoskeletal dynamics affects cell cycle progression. Previous studies have shown that Rho signalling both potentiates focal adhesion assembly and relieves p27-dependent cell cycle arrest (Hotchin & Hall, 1995; Mammoto *et al*, 2004). However, following 4OHT treatment, the proliferation of *Mrtfa^-/-^*;*Mrtfb^fl/fl^* cells was not restored upon expression of constitutively active RhoA^G14V^ (Figure S7A, S7B). We therefore investigated whether interference with cytoskeletal dynamics in wildtype MEFs could affect cell cycle progression. We focussed on cell contractility, which is required for progression through the G1/S checkpoint in endothelial cells (Huang *et al*., 1998). The Rho kinases (ROCKs) are important regulators of cell contractility, controlling both F-actin assembly and myosin activity (reviewed by Rath & Olson, 2012), and ROCK inhibition was previously reported to lead to both cytokinesis defects and cellular senescence (Kumper *et al*, 2016).

Treatment of wildtype MEFs with the pan-ROCK inhibitor H1152 reduced actin stress fibres and focal adhesions, as did serum-deprivation, but did not induce the cell rounding seen in *Mrtfab^-/-^*MEFs (Figure S7C). Nevertheless, H1152 reduced proliferation of immortalised WT MEFs to a level comparable to that of *Mrtfab^-/-^* MEFs (Figure 7D-F). ROCK-inhibited wildtype MEFs exhibited similar changes in p27, CDK1 and CKS2 expression to *Mrtfab^-/-^*MEFs and serum-deprived wildtype MEFs (Figure 7G), as well as increased SA-βGal activity, and elevated transcription of SASP markers (Figure S7E, S7F). In contrast to *Mrtfab^-/-^*MEFs, however, H1152 treatment of wildtype MEFs induced significant numbers of multinucleated cells (Figure S7C,D), as expected given the role of the ROCKs in cytokinesis (Green *et al*, 2012). Treatment with the Myosin II inhibitor blebbistatin also inhibited proliferation of WT MEFs (Figure 7D, 7H). Blebbistatin-treated wildtype MEFs exhibited similar cytoskeletal changes to ROCK-inhibited cells (Figure S7C), and blebbistatin treatment induced similar effects on cell cycle marker expression to H1152 treatment or serum deprivation of wild type cells, or MRTF inactivation (Figure 7G). Finally, we directly examined contractility in *Mrtfab^-/-^*MEFs using traction force microscopy. *Mrtfab^-/-^* MEFs exhibited reduced contractility, exerting 70% less force per unit area than wildtype MEFs (Figure 7I). Taken together, these results suggest that the proliferation defect observed in *Mrtfab^-/-^* MEFs is likely due to intrinsic defects in actin dynamics, and that this at least in part reflects reduced contractility.

## DISCUSSION

In this study we have shown that the MRTF-SRF axis plays a significant role in cell proliferation in fibroblast and epithelial cells. Inactivation of SRF, or of MRTF-B in fibroblasts lacking MRTF-A, induces a profound proliferative defect, with decreased levels of cyclin D, CDK1 and CKS2, increased amounts of the CDK inhibitor p27, elevated neutral β-galactosidase activity, and expression of numerous SASP markers. Similar phenotypes are also seen in wildtype cells upon serum-starvation, or upon inhibition of the ROCKs or actomyosin contractility. MRTF-SRF inactivation results in multiple cytoskeletal defects, including substantially reduced contractility. MRTF-A and MRTF-B function redundantly to promote cytoskeletal integrity and cell proliferation. Our findings suggest that MRTF-SRF dependent cytoskeletal dynamics, particularly contractility, play a pivotal role in generating the pro-proliferative signal provided by cell-substate adhesion in adherent cells.

Several aspects of the *Mrtfab^-/-^* phenotype are seen in senescent cells, but we do not consider MRTF-null cells to exhibit classical senescence. The stringent definition of senescence has been hampered by the lack of well-defined and specific markers (Munoz-Espin & Serrano, 2014). MRTF-null MEFs exhibit a SASP-like secretory phenotype, and can inhibit proliferation of co-cultured wildtype cells. However, SASP-like phenotypes have also been reported in quiescent cells (Anwar *et al*, 2018), and recent studies suggest that the SASP and SA-βgal reflect duration of cell cycle withdrawal rather than a distinct non-proliferative state (Ashraf *et al*, 2023). Irreversibility has been conventionally been considered a hallmark of senescence (Campisi & d’Adda di Fagagna, 2007), but we found that all MRTF-null MEF phenotypes can be fully reversed by MRTF-A re-expression. Finally, although we found that *Mrtfab^-/-^* MEFs apparently exhibit increased cell plasticity, a phenotype associated with therapy-induced cancer cell senescence (Milanovic *et al*, 2018), MRTF-SRF activity has also been previously reported to suppress plasticity in non-senescent cells (Hu *et al*., 2019).

Previous studies of the MRTFs and SRF have led to the view that the pathway affects cell motile and adhesive properties rather than cell proliferation. SRF-null embryos proliferate normally up to E6 but fail to gastrulate (Arsenian *et al*., 1998; Schratt *et al*., 2001), and SRF-null neuroblasts form in normal numbers but are unable to migrate to the olfactory bulb (Alberti *et al*., 2005). Inactivation of SRF or the MRTFs does not appreciably impair the early proliferative stages of T cell differentiation or activation (Fleige *et al*, 2007; Maurice *et al*., 2024; Mylona *et al*., 2011). However, in both human and mouse, smooth muscle cells that lack SRF exhibit an increased propensity to enter senescence (Angstenberger *et al*., 2007; Hengst *et al*, 1994; Werth *et al*., 2010), while MRTF-SRF signalling suppresses oncogene-induced senescence (OIS) in cancer cells lacking the tumor suppressor DLC1, a RhoGAP (Hampl *et al*., 2013; Hermanns *et al*., 2017). Although further work is needed to elucidate the basis for these context-dependent differences, our data show that MRTF-SRF signalling is likely to play a more general role in proliferation than previously thought.

Our findings show that MRTF inactivation induces a quiescence-like state under conditions that permit efficient cell cycle progression in wildtype cells. Elevated p27 levels are characteristic of cell cycle inhibition induced in various cell contexts by non-genotoxic cues including serum deprivation (Coats *et al*, 1996), TGF-β treatment or contact inhibition (Polyak *et al*, 1994), restriction of cell spreading (Chen *et al*., 1997; Huang *et al*., 1998), low matrix stiffness (Klein *et al*., 2009), or lovastatin treatment (Hengst *et al*., 1994; Hengst & Reed, 1996). Many of these directly affect activation of the MRTF-SRF signalling pathway, or impinge on cytoskeletal dynamics. At least in some settings, arrest-induced p27 accumulation, which is Rho-dependent, reflects decreased Skp2 levels (Huang *et al*., 1998; Mammoto *et al*., 2004). Decreased Skp2 levels are also seen in SRF-null VSM cells (Werth *et al*., 2010). Both Skp2 protein and transcript levels were slightly decreased in our MRTF- null MEFs, and we note that in breast cancer cells, Skp2 expression also responds to mechanical cues via the YAP pathway (Jang *et al*, 2017).

The CDK inhibitors p27 and p21 inhibit CDKs through a common mechanism (Polyak *et al*., 1994), and p27 and p57 share functions in proliferation control (Matsumoto *et al*., 2011; Susaki *et al*., 2009). We found that in contrast to immortalised *Mrtfab^-/-^*MEFs, *Mrtfab^-/-^* primary kidney fibroblasts and tracheal epithelial cells exhibited elevated p21 levels, with correspondingly reduced p27 levels. While this suggests that p27 or other CIP/KIP family proteins might function redundantly to mediate the proliferation defect, we found that inactivation of neither p27 nor all three CIP/KIP proteins was sufficient to restore proliferation to *Mrtfab^-/-^* MEFs. This finding extends previous studies showing that MEFs lacking individual CIP/KIP family members retain growth control by contact inhibition and serum deprivation (Deng *et al*, 1995; Nakayama *et al*., 1996; Takahashi *et al*, 2000). We therefore suggest that CKI elevation upon MRTF inactivation is a consequence rather than a cause of proliferation arrest in MEFs.

We found culture of wildtype MEFs on limited areas of adhesion also reduced cell proliferation to a comparable extent to unconfined *Mrtfab^-/-^* cells. Such confinement also increases cellular G-actin levels, which would be expected to decrease MRTF activity (R Fedoryshchak *et al*, manuscript in preparation). In contrast proliferation was not affected when wildtype cells are allowed to spread under conditions of limited adhesion area in agreement with previous findings (Chen *et al*., 1997), and in this case G-actin levels are unaffected. We therefore speculate that primary driver for confinement-mediated impairment of cell proliferation may be altered actin dynamics. We note, however, that the greatly reduced proliferation of *Mrtfab^-/-^* MEFs can also be further inhibited by confinement, suggesting other mechano-responsive pathways promote proliferation. One such pathway might be the mechano-responsive YAP/TAZ system (Dupont *et al*, 2011).

The quiescence-like state associated with MRTF inactivation can also be induced in wildtype MEFs by treatment with the ROCK1/ROCK2 inhibitor H1152 or the myosin inhibitor blebbistatin. Both ROCKs and myosin are critical for cell contractility, and MRTF-null MEFs exhibit substantially impaired contractility. Moreover, classical studies have shown that F-actin integrity and cell contractility are required for progression through the G1 restriction point (Huang *et al*., 1998; Iwig *et al*, 1995; Kumper *et al*., 2016; Reshetnikova *et al*, 2000). How might this affect cell progression? Two non-mutually exclusive mechanisms can be envisaged. First, MRTF inactivation might directly limit contractility-dependent changes in cell morphology required for cell cycle progression. For example, during G2/M, ROCK- and myosin-dependent contractility promotes mitotic cell rounding by multiple mechanisms (reviewed by Ramkumar & Baum, 2016) including focal adhesion disassembly and the assembly of rigid cortical actomyosin (Jones *et al*, 2018; Matthews *et al*, 2012; Ramanathan *et al*, 2015), and the actomyosin contractile ring at cytokinesis (Green *et al*., 2012). Moreover, in pseudo-stratified epithelia such as the tracheal epithelial cells studied here actomyosin contractility is essential for the nuclear movements required for mitosis (reviewed by Meyer *et al*, 2011). A second possibility is that MRTF inactivation limits the generation of pro-proliferative signals from the cytoskeleton. Mechanical cues, including cell adhesion and spreading (Chen *et al*., 1997; Huang *et al*., 1998; Mammoto *et al*., 2004), and matrix compliance (Klein *et al*., 2009; Mih *et al*, 2011) promote cell cycle progression in anchorage-dependent cells. Focal adhesions, whose assembly is MRTF-SRF dependent in our cells, would be a prime candidates to generate such signals (for review see Burridge & Chrzanowska-Wodnicka, 1996; Kamranvar *et al*., 2022).

MRTF-SRF target genes include both components and regulators of the actin cytoskeleton, including the actins themselves (Schratt, 2002 #21, reviewed by Olson & Nordheim, 2010). We propose that the contractily deficits underpinning defective proliferation of MRTF-null cells reflect reduced expression of multiple cytoskeletal MRTF-SRF target genes, although it remains possible that MRTF-SRF also directly controls expression of cell cycle regulators. G-actin plays a critical role in regulation of MRTF activity, and in at least some contexts overexpression of cytoplasmic actin can rescue MRTF-dependent phenotypes (Maurice *et al*., 2024; Salvany *et al*, 2014). Intriguingly, in MEFs the inactivation of cytoplasmic actin genes results in defective proliferation (Bunnell *et al*, 2011; Patrinostro *et al*, 2017; Tondeleir *et al*, 2012). The proliferative and cytoskeletal phenotypes of MRTF-null cells may thus at root reflect deficient actin expression, and we are currently testing this idea.

We have shown that MRTF-SRF activity is required for proliferation of anchorage dependent fibroblasts and epithelial cells, most likely because MRTF-SRF dependent cytoskeletal dynamics and contractility allow adherent cells to execute the morphological changes required for cell division. However, many untransformed cell types also proliferate independently of adhesion, and anchorage-independent proliferation is also a hallmark of oncogenic transformation. In principle oncogenic signals could simply substitute for pro-proliferative signals generated in nontransformed cells by substrate adhesion. According to this view, proliferation of transformed cells might occur independently of MRTF-SRF activity, providing that constraints on cytoskeletal dynamics are not limiting, which might well be the case for non-adherent cells. Our future work will focus on the relation between transformation, MRTF-SRF activity, and the cytoskeleton.

## Supporting information

TABLE 1 - RNAseq data

TABLE 2 - GSEA and GO analysis

## ACKNOWLEDGEMENTS

We thank Patrick Costello, lab members, Michael Way, and Francesco Gualdrini for helpful discussions, and Gerard Evan, Michael Way and Paul Nurse for helpful comments on the manuscript. We thank Stefania Crotta and Andreas Wack for help with tracheal epithelial cell culture, the Sahai lab for the Fast-Fucci plasmid (Addgene #86849), the Way lab for pLVX-mCherry plasmid (Addgene #180646). Mike Howell and Ming Jiang from the Crick High-throughput screening STP for help with generation of the Cas9 cell line, Robert Goldstone and Deb Jackson in the Crick Advanced sequencing STP for library preparation and sequencing, and Donald Bell and Matthew Renshaw from the Crick Advanced Light Microscopy STP for guidance and assistance using microscopes and slide scanner.

This work was supported by the Francis Crick Institute which receives its core funding from Cancer Research UK (CC2102), the UK Medical Research Council (CC2102), and the Wellcome Trust (CC2102). This research was funded in whole, or in part, by the Wellcome Trust CC2102. For the purpose of Open Access, the author has applied a CC BY public copyright licence to any Author Accepted Manuscript version arising from this submission. The authors declare that they have no conflicts of interest.

## AUTHOR CONTRIBUTIONS

JN designed and conducted experiments, drew figures and drafted the manuscript. MB-J assisted with derivation of tissue-specific fibroblasts, and cell adhesion, contractility experiments. NP conducted gene inactivation and re-expression experiments. JD cultured tracheal epithelial cells, established conditions for conditional MRTF inactivation, and conducted experiments. SB conducted initial characterisation of MRTF- and SRF- depleted cultures, and established the similarity between the ROCK-inhibited and MRTF-null phenotype. RT established the project, designed and interpreted experiments, and with JN wrote the paper with input from the other authors.

Conceptualization RT, JN, SB

Data curation JN

Formal Analysis JN

Funding acquistion RT

Investigation JN, MB-J, NCP, JD, SB

Supervision RT, JN

Writing – original JN, RT

Review and editing JN, RT, MB-J, NP, JD

## SUPPLEMENTARY FIGURE LEGENDS

**Figure S1:**
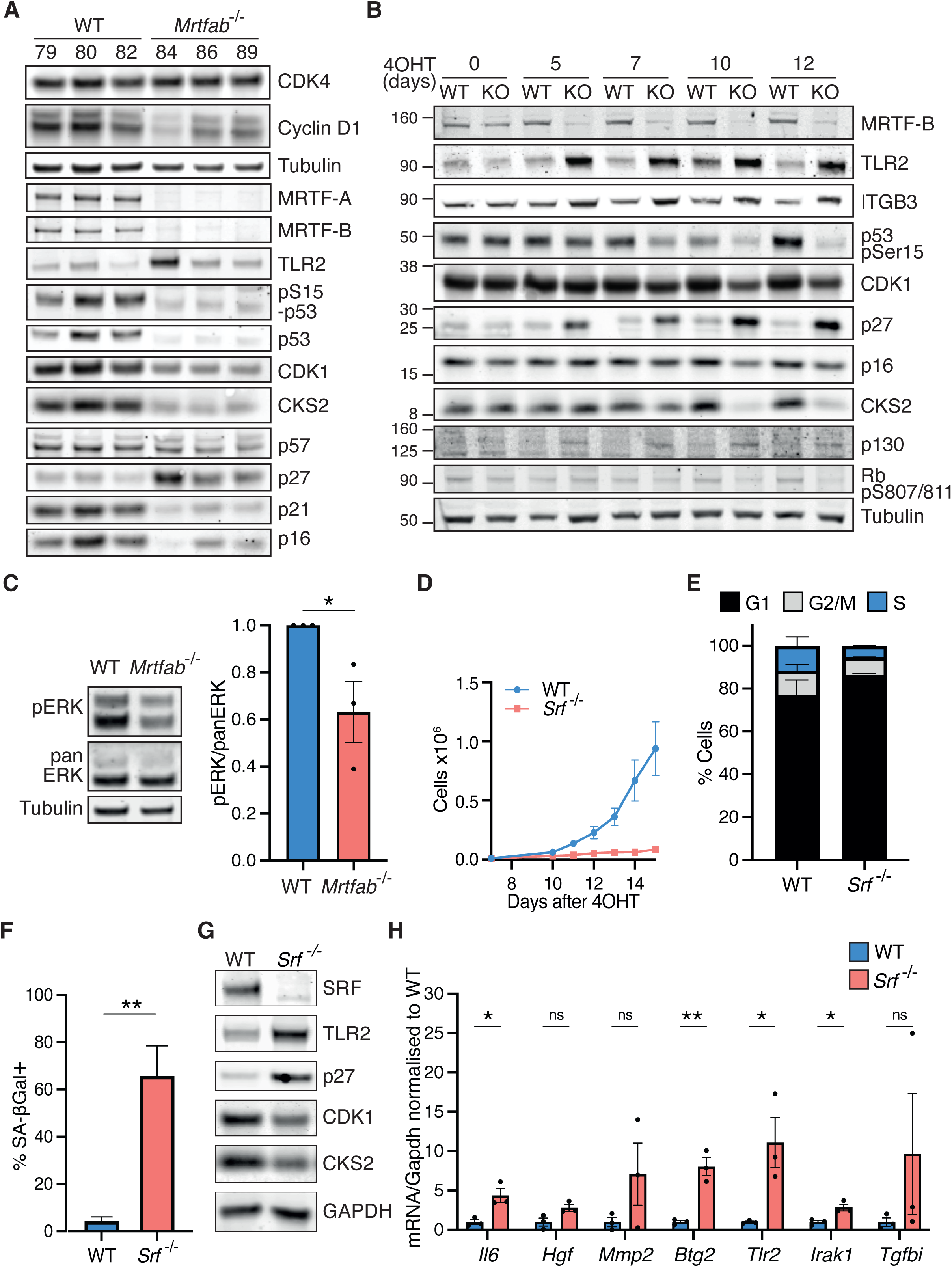
Defective proliferation of MRTF- or SRF-null MEFs. A) Immunoblot analysis of cell cycle and senescence markers in each pool of WT or *Mrtfab*^-/-^ MEFs at day 12 after 4OHT. B) Immunoblotting of WT (pool 80) and *Mrtfab*^-/-^ (pool 86) MEFs at the indicated times s after 4OHT. Similar results were obtained with the other pools. C) ERK activity in day 12 WT and *Mrtfab*^-/-^ MEFs. Left. representative immunoblot of pERK and panERK (pools 80 and 86). Right, quantification showing mean of the three independent pools of each genotype. Results are normalised to WT. Statistical analysis, unpaired t-test. *, p<0.05. D) Pools of MEFs derived from WT or *Srf*^-/-^ MEFs (three pools per genotype, each derived from one embryo) were treated with 4OHT and growth curves determined as in Figure 1A. For each pool, cell counts at each time point are the mean of three technical replicates. Data points are the mean cell counts of the three independent cell pools ± SEM. A representative experiment is shown. E) Cell cycle distribution of WT or *Srf*^-/-^ MEFs on day 12 after 4OHT evaluated by PI staining. F) SA-βgal staining of WT and *Srf*^-/-^ MEFs on day 12 after 4OHT. Data are mean of the three independent pools (>100 cells scored per pool) ± SEM. Statistical analysis, unpaired t-test. **, p<0.01. G) Immunoblotting of WT or *Srf*^-/-^ MEFs on day 12 after 4OHT. Data are from pools MEF4 and Bis1. Similar results were obtained with the other pools. H) RT-qPCR of mRNA levels of SASP factors in day 12 WT or *Mrtfab*^-/-^ MEFs after 4OHT. Data are mean of the three independent pools ± SEM; significance, unpaired t-test. ns, not significant; *, p<0.05; **, p<0.01.

**Figure S2:**
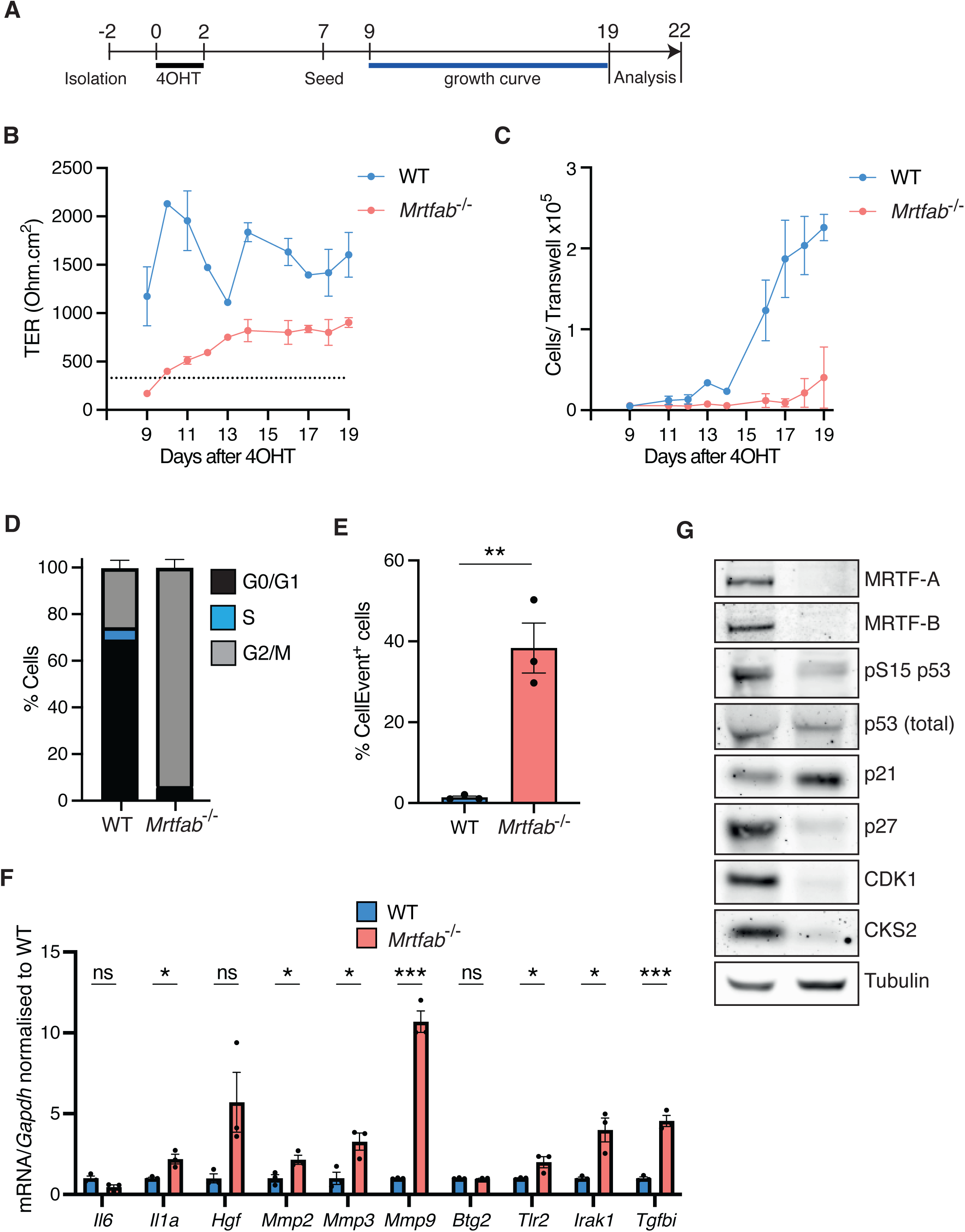
Defective proliferation of MRTF-null primary tracheal epithelial cells. A) Experimental protocol. Mouse tracheal epithelial progenitor cells were isolated, treated with 4OHT, and seeded on collagen/fibronectin coated transwell inserts (10,000 cells per insert). B) Epithelial barrier function, evaluated by measurement of transepithelial resistance (TER). Data are mean of three independent experiments ± SEM. For the different experiments 4, 3-10 or 6-21 technical replicates were quantified. Dotted line, resistance of confluent cell sheet. C) Cells from (B) were trypsinised and counted over the course of 12 days. D) Cell cycle distribution analysis of day 20 BrdU-labelled / PI-stained epithelial cells. Data is from a single experiment with 3 technical replicates ± SEM. E) Proportion of Cell Event Senescence Green positive epithelial progenitor cells as measured by flow cytometry. Data are mean of three independent experiments on on days 19 to 22 after 4OHT, ± SEM. Significance, unpaired t-test; p<0.01. F) RT-qPCR of mRNA levels of SASP factors, normalised to WT cells on day 21 after 4OHT. Data is from a single experiment with 3 technical replicates, ± SEM. Significance, multiple unpaired t-test; ns, not significant; *, p<0.05; ***, p<0.001. G) Immunoblotting of epithelial progenitor cells on day 19 after 4OHT treatment. Representative of three independent experiments.

**Figure S3:**
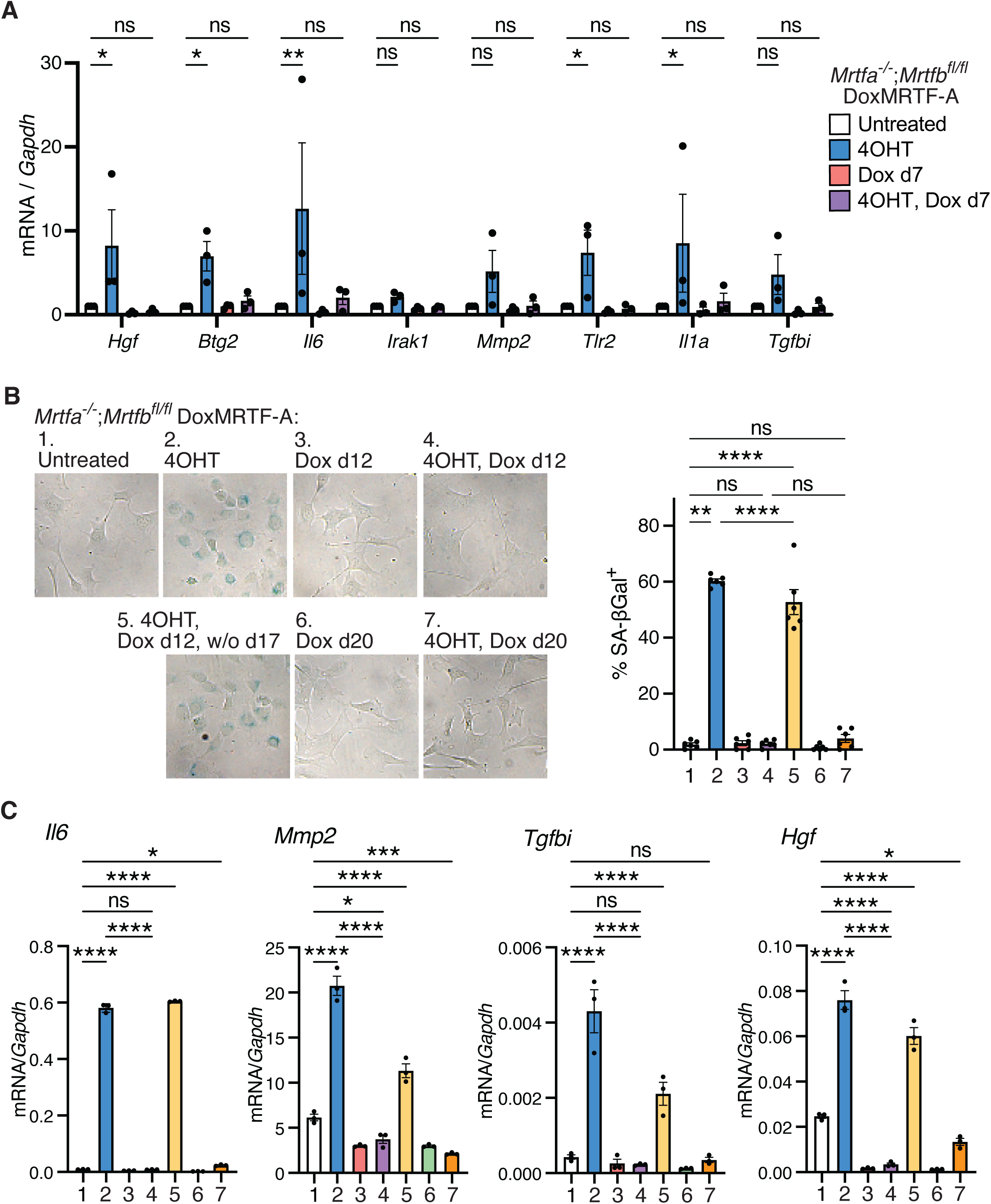
MRTF-A re-expression reverses the proliferative and cytoskeletal defects of *Mrtfab*^-/-^ MEFs. A) RT-qPCR of SASP markers at day 12 in the three independent *Mrtfab^-/-^* DoxMRTF-A cell lines treated as in Figure 3A. Data are mean of the three independent lines ± SEM, normalised to untreated cells. Statistical analysis, two-way ANOVA with correction for multiple comparisons. ns, not significant; *, p<0.05; **, p<0.01. B) Left, representative images of SA-βGal staining at day 25 in *Mrtfab^-/-^* DoxMRTF-A cells 1F2 treated as in Figure 3D. Right, quantification; conditions as at left. Data are mean of 6 technical replicates ± SEM. Statistical analysis one-way ANOVA with correction for multiple comparisons; ns, not significant; **, p<0.01; ****, p<0.0001. C) RT-qPCR of SASP markers at day 25 in *Mrtfab^-/-^* DoxMRTF-A 1F2 cells treated as in (B). Data are mean of technical replicates ± SEM. Similar results were obtained with 1F5 cells. Statistical analysis, two-way ANOVA with correction for multiple comparisons; ns, not significant; *, p<0.05; ***, p<0.001; ****, p<0.0001.

**Figure S4:**
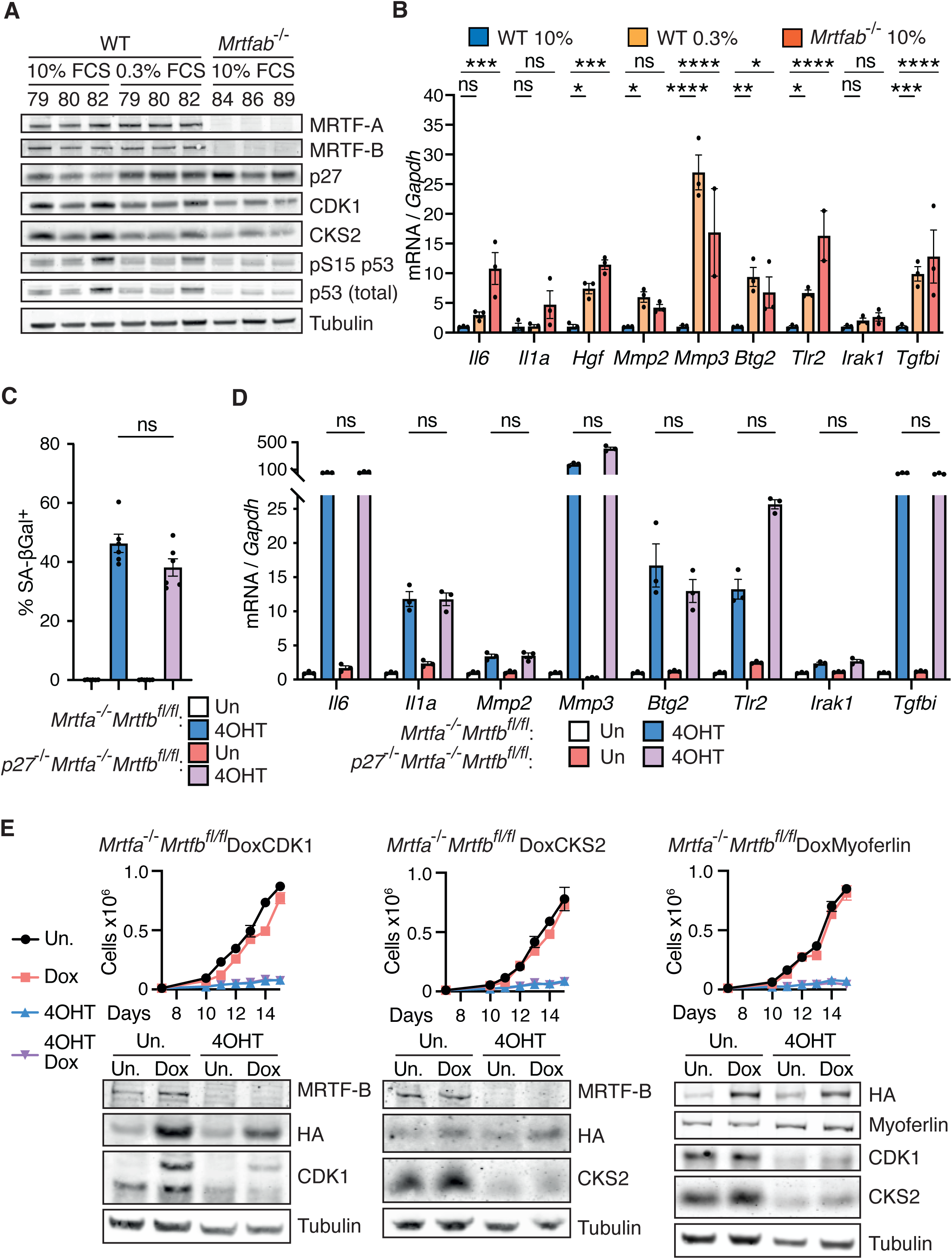
The MRTF-null phenotype exhibits features of quiescence. A) Cells were treated as in Figure 4A but with medium was changed to 0.3% FCS on day 10 *ie* 2 days serum deprivation. B) RT-qPCR of SASP mRNA levels in cells treated as in Figure 4A. Data are mean ±SEM of three pools of each genotype. Statistical analysis used two-way ANOVA with multiple comparisons using Fisher’s LSD test: ns, not significant; *, p<0.05; **, p<0.01; ***, p<0.001;****, p<0.0001. C) Line 3A2 p27^-/-^*Mrtfa*^-/-^*Mrtfb*^fl/fl^ MEFs treated as in Figure 4F were stained for SA-βGal activity on day 12. Data are mean ± SEM of 6 technical replicates. Statistical analysis was by unpaired t-test: ns, not significant. D) SASP marker expression in line 3A2 p27^-/-^*Mrtfa*^-/-^*Mrtfb*^fl/fl^ MEFs treated as in Figure 4F were analysed by RT-qPCR on day 12. Data are mean ± SEM of three technical replicates. Statistical analysis was by multiple Mann-Whitney tests. ns, not significant. E) Pool 86 *Mrtfa*^-/-^*Mrtfb*^fl/fl^*Rosa26*^Tam-Cre^ cells were infected with lentiviruses encoding doxycycline-inducible HA-tagged CDK1 (left), CKS2 (centre) or Myoferlin (right). Following puromycin selection, the resulting cell pools were treated or not with 4OHT and/or doxycycline. Top panels, cell proliferation from days 10 to 15. Data are means of three independent experiments ± SEM. Bottom panels, cell cycle marker expression at day 12.

**Figure S5:**
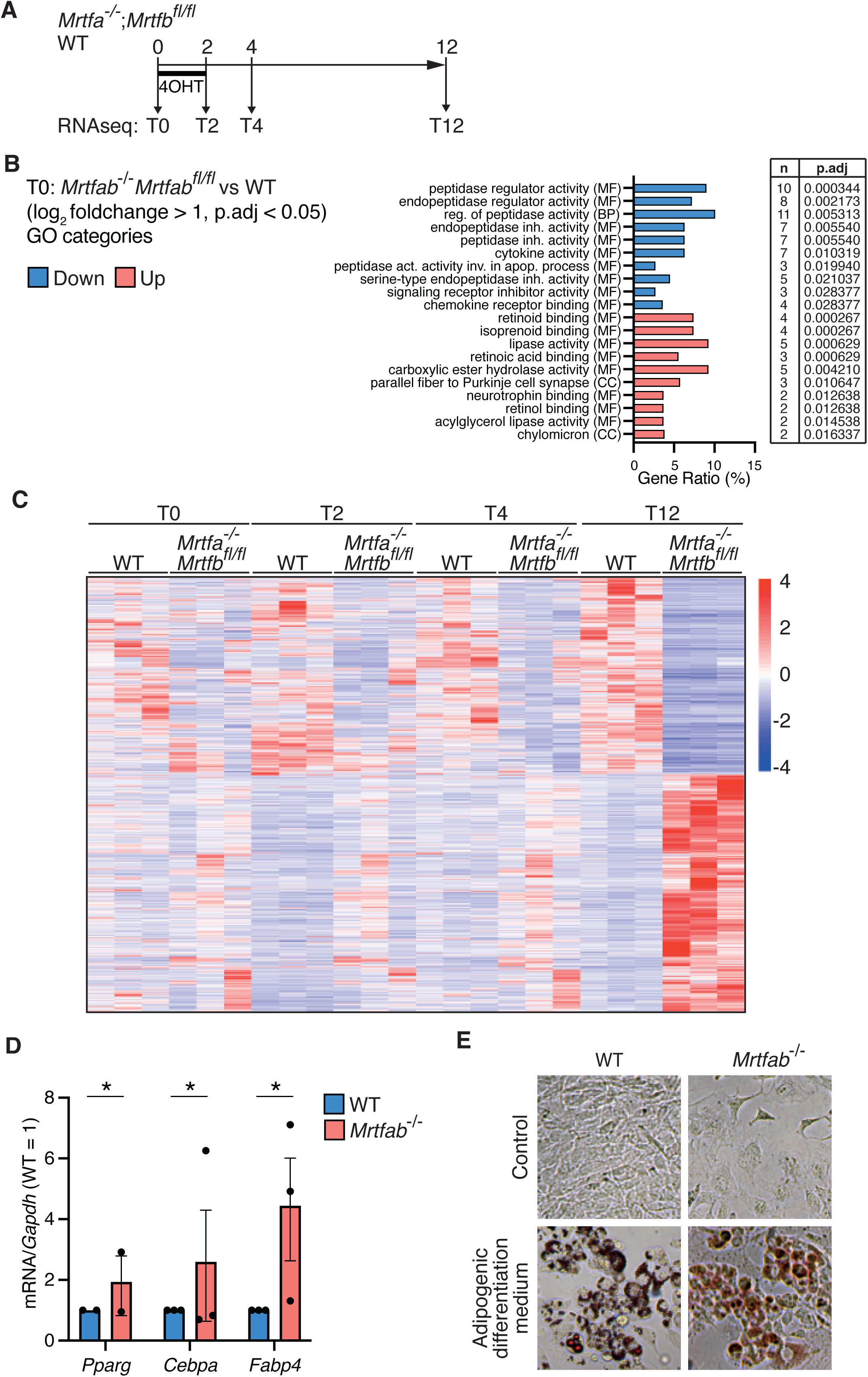
MRTF inactivation results in a decrease in actin-related genes consistent with previous identified MRTF target genes. A) Experimental protocol. WT or *Mrtfa^-/-^*;*Mrtfb^fl/fl^* MEFs on were treated with 4OHT on day 0 and RNA collected for RNAseq analysis on day 0, 2, 4, and 12 (samples T0, T2, T4 and T12 respectively). B) Gene Ontology (GO) analysis of 189 genes significantly up- or downregulated (>2-fold change, Padj <0.05) in *Mrtfa*^-/-^*Mrtfb^fl/fl^*T0 pools compared with wildtype MEF pools. The top 10 GO terms (BP: Biological Process, MF: Molecular Function, CC: Cellular component) the Benjamini-Hochberg adjusted p-value for each set. C) WT MEF pools 79, 80, 82; and *Mrtfa*^-/-^*Mrtfb^fl/fl^* MEF pools 84, 86, 89 were treated or not with 4OHT and cultured for 0, 2, 4, or 12 days before RNAseq analysis (T0, T2, T4, T12 samples, see Figure 5A). Plot shows z-scores of genes showing a >2-fold change with Padj <0.05 for each pool in all samples, ordered according to unsupervised clustering of the T12 data. T0 and T12 data also shown in Figure 5A. D) RT-qPCR analysis of adipogenic markers *Pparg*, *Cebpa*, and *Fabp4*. Data are mean of three independent experiments done in triplicate ± SEM. Statistical analysis by two-way ANOVA; p<0.05. E) Oil Red O staining of WT or *Mrtfab*^-/-^ MEFs cultured in regular or adipogenic differentiation medium (with 0.5mM IBMX, 1µM dexamethasone, 10µM insulin) for 20 days. Data are representative of two independent experiments.

**Figure S6.**
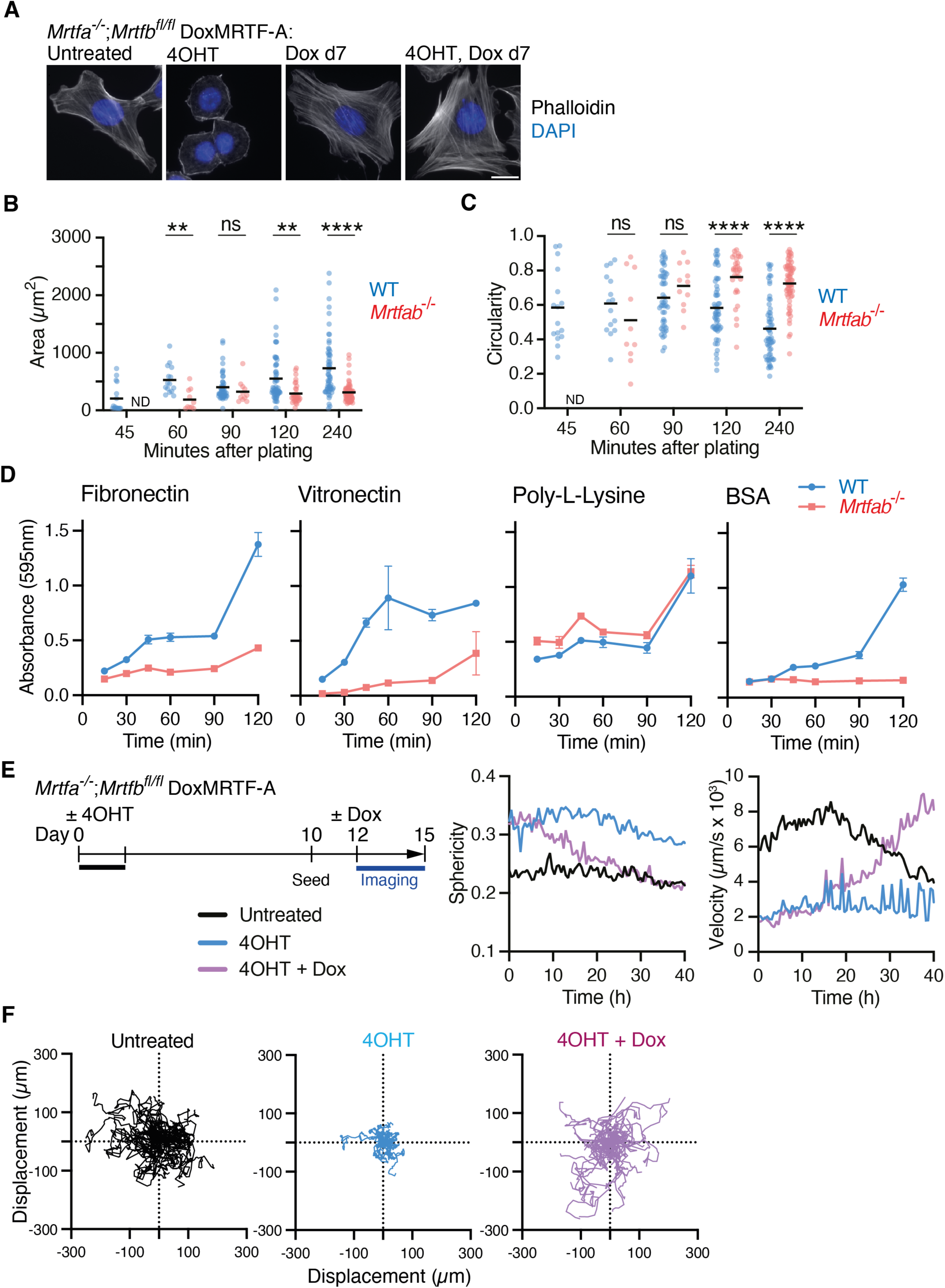
Cytoskeletal defects in *Mrtfab*^-/-^ MEFs. A) Cell morphology at day 12 in line 1F2 *Mrtfab^-/-^* DoxMRTF-A cells treated as in Figure 3A. F-actin is stained using phalloidin. Images are representative images of 2 biological replicates (1F2 and 1F5 cell lines). Scale bar, 20µm. B) WT or *Mrtfab*^-/-^ MEFs (pools 80 and 86 respectively) were seeded and stained with CellMask Orange followed by fixation at indicated times after seeding. Cell areas were measured (>10 cells per timepoint); mean is indicated by the line. Statistical analysis, unpaired t-test; ND, not determined; ns, not significant; **, p<0.01; ****, p<0.0001. C) Circularity of cells from (B). Analysis as in (B). D) Adhesion assay on fibronectin, vitronectin, Poly-L-Lysine, or BSA. 50,000 cells pool 80 WT or pool 86 *Mrtfab*^-/-^ MEFs were seeded per assay, stained with crystal violet at the indicated times, and adherent numbers quantified by absorbance at 595nm. Data are mean of three technical replicates ± SEM, and are representative of 5 independent experiments. E) Protocol for phasefocus experiment. *Mrtfa^-/-^*;*Mrtfb^fl/fl^* DoxMRTF-A cells (line 1F2) were treated or not with 4OHT and/or subsequently with doxycycline, and imaged for the next three days at 2 images / h (see Videos 3-5) and mean sphericity instantaneous velocity determined. Velocity of the untreated cells decreases at later times as cell migration becomes compromised by confluence. Note that upon Dox treatment GFP+ cells, re-expressing MRTF-A, accumulate slowly over a period of 10-15h in 4OHT-treated *Mrtfa^-/-^Mrtfb^fl/fl^* cells (see Video 5). F) Displacement plots of 1F2 cells treated as in (E). Cells (were monitored until division, exit of frame of view, or end of the video and 50 cell tracks plotted for each culture. Similar results were obtained with line 1F5 and 1B2.

**Figure S7:**
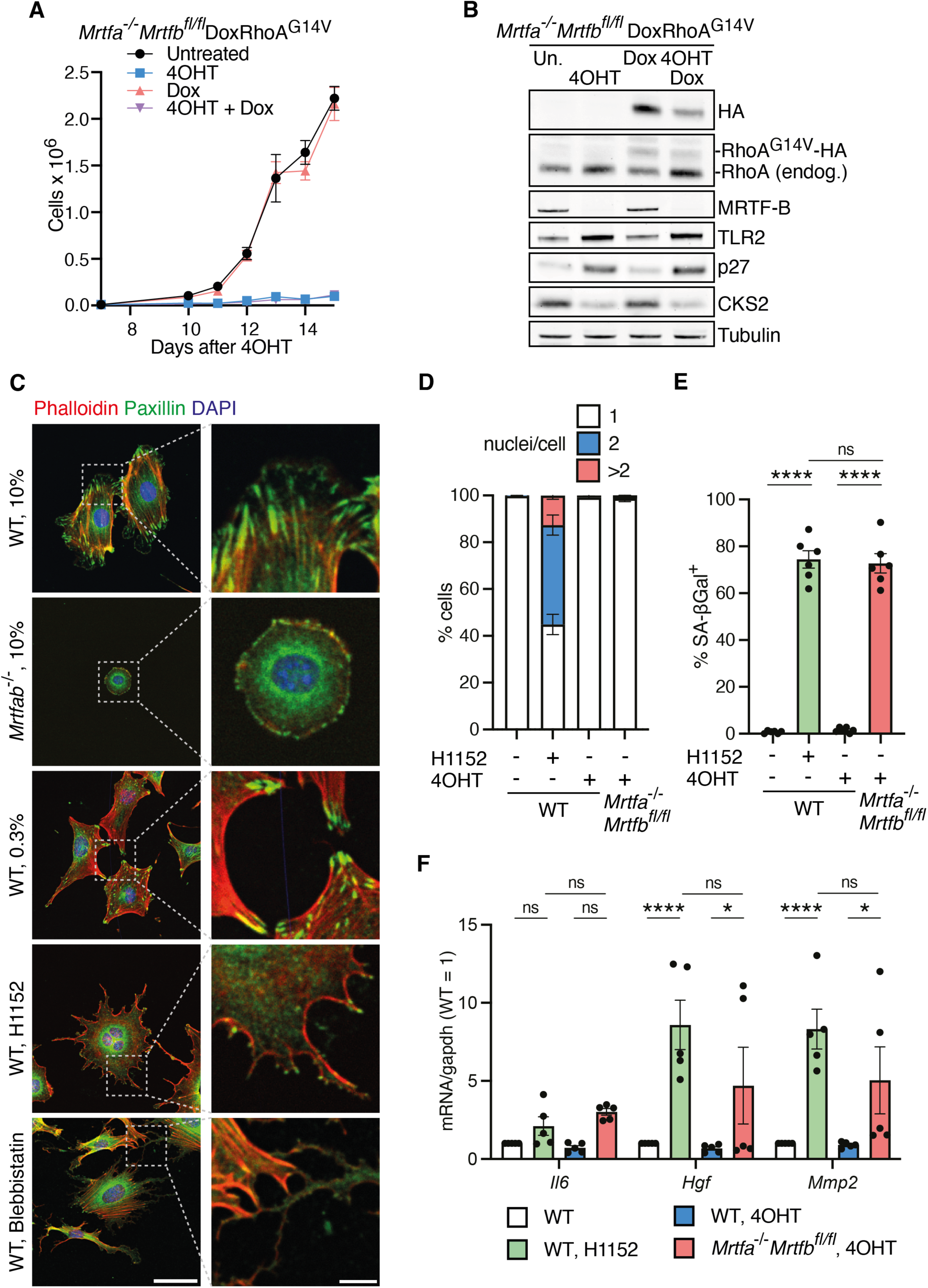
Inhibition of contractility in wildtype cells induces a phenoptype similar to MRTF inactivation. A) Pool 86 *Mrtfa*^-/-^*Mrtfb*^fl/fl^*Rosa26*^Tam-Cre^ MEFs were infected with lentivirus pCW- RhoA^G14V^-HA carrying dox-inducible constitutively active RhoA. Two days after plating, cells were treated or not with 4OHT and/or doxycycline. Cells were counted on day 10-15 after 4OHT. Data are mean ± SEM from 3 independent experiments. B) Representative immunoblot of pool 86 *Mrtfa*^-/-^*Mrtfb*^fl/fl^DoxRhoA^G14V^ cells from (A) on day 12 after 4OHT treatment. C) Immunofluorescence staining of paxillin (focal adhesions), phalloidin (F-actin) and DAPI (DNA) in pool 80 WT or pool 86 *Mrtfab*^-/-^ MEFs treated as in Figure 7D. Scale bar = 50µm (left) and 10µm (right). Representative of 3 independent experiments. D) Nuclear counts in pool 80 WT or pool 86 *Mrtfab*^-/-^ MEFs on day 12 after 4OHT and/or H1152 treatment as in Figure 7D. >10 cells analysed per replicate sample Data mean ± SEM of 6 replicates. E) SA-βgal staining of pool 80 WT or pool 86 *Mrtfab*^-/-^ MEFs on day 12 after 4OHT and/or H1152 treatment as in Figure 7D. Data are mean ± SEM of three individual experiments, >100 cells scored per sample, n=3; significance, unpaired t-test; ns, not significant; ****, p<0.0001. F) RT-qPCR of mRNA levels of SASP factors in pool 80 WT or pool 86 *Mrtfab*^-/-^ MEFs on day 12 after 4OHT and/or H1152 treatment as in Figure 7D. Data are mean of 5 independent experiments ± SEM; significance, unpaired t-test; ns, not significant; *, p<0.05; ****, p<0.0001.

## TABLE LEGENDS

**TABLE 1. Differential gene expression upon MRTF inactivation in MEFs**

WT or *Mrtfa^-/-^*;*Mrtfb^fl/fl^* MEFs were treated with 4OHT on day 0, and RNA collected for RNAseq analysis on day 0, 2, 4, and 12. Differential gene expression analysis was carried out (samples T0, T2, T4 and T12 respectively). Each table presents differential gene expression analyses between WT and *Mrtfa^-/-^Mrtfb^fl/fl^* (MRTFdKO) MEFs at each time point, ranked from maximum fold-downregulation to maximum fold-upregulation in MRTFdKO MEFs compared with wildtype MEFs. Columns are: ENSEMBL, ensemble gene ID; baseMean, average normalised expression across samples; log2FoldChange, log_2_-transformed fold change between conditions; lfcSE, standard error of the log_2_ fold change; pvalue, raw p-value from Wald test; padj, p-value adjusted for multiple testing using Benjamini-Hochberg (BH); SYMBOL, gene symbol; ENTREZID, Entrez gene ID; −log10(pvalue), negative log_10_-transformed adjusted p-value for volcano plots.

**TABLE 2. GSEA and GO analysis of genes downregulated at day 0 or 12 after MRTF-inactivation**

Sheet 1: Gene Set Enrichment Analysis (GSEA) table at T12 showing significant gene sets of the Hallmark category. Analysed in R using GSEA() from clusterprofiler with Hallmark gene sets from msigdbr. Columns are: ID, Hallmark identifier of gene set; setSize, number of genes from the gene set present in the data; enrichmentScore, running-sum statistic of degree of enrichment; NES, Normalised Enrichment Score – enrichment score normalised by gene set; pvalue, raw p-value from permutation test of enrichment; p.adjust, p-value adjusted for multiple comparisons using Benjamini-Hochberg (BH); qvalue, estimated false discovery rate (FDR); rank, position of peak enrichment score in the ranked gene list; leading_edge, summary of genes driving enrichment; core_enrichment, gene symbols in leading edge subset.

Sheet 2: Gene Ontology (GO) analysis table at T12 showing significant upregulated gene ontology gene lists. Analysed using enrichGO() from clusterprofiler with ont = “ALL”. Columns are: ONTOLOGY, category; ID, GO term identifier; Description, name of gene ontology term; GeneRatio, proportion of input genes in the GO term; BgRatio, proportion of background genes in the GO term; pvalue, raw p-value from hypergeometric test; p.adjust, p-value adjusted for multiple comparisons using Benjamini-Hochberg (BH); qvalue, estimated false discovery rate (FDR); geneID, input genes annotated to the GO term; Count, number of input genes annotated to the GO term.

Sheet 3: As sheet 2, but for significantly downregulated genes at T12. Sheet 4: As sheet 2, but for significantly upregulated genes at T0.

Sheet 5: As sheet 2, but for significantly downregulated genes at T0.

## SUPPLEMENTARY TABLES

**TABLE S1. Oligonucleotides and guides**

**TABLE S2. Antibodies.**

## VIDEO LEGENDS

**Videos 1 and 2. Timelapse microscopy of wildtype and *Mrtfab*^-/-^ MEFs.**

Cells were filmed over a 96h period, with 1 frame per 30 minutes.

**Videos 3-5. MRTF-A reexpression restores morphology, motility, and proliferation.** *Mrtfa^-/-^Mrtfb^fl/fl^* DoxMRTF-A cells were treated as in Figure 5E and analysed by Phasefocus livecyte microscopy at day 12 for 40h at 2 images/h.

Videos 3a, 4a, 5a, 6a: GFP images. Videos 3b, 4b, 5b, 6b: phase contrast images. Video 3a, 3b: untreated cells.

Videos 4a,4b: cells treated with 4OHT.

Videos 5a, 5b: 4OHT-treated cells at day 12 treated with Dox 1h prior to imaging.

Velocity of the untreated cells decreases at later times as cells become confluent, which was not reached by the Dox-treated cells (compare videos 3,5).

Note that upon Dox treatment GFP+ cells, re-expressing MRTF-A, accumulate slowly over a period of 10-15h in 4OHT-treated *Mrtfa^-/-^Mrtfb^fl/fl^* cells (Videos 5a).

## Notes

https://www.ncbi.nlm.nih.gov/geo/query/acc.cgi?acc=GSE298922

### Competing Interest Statement

The authors have declared no competing interest.

